# Structure-guided aminoacylation and assembly of chimeric RNAs

**DOI:** 10.1101/2024.03.02.583109

**Authors:** Aleksandar Radakovic, Anna Lewicka, Marco Todisco, Harry R. M. Aitken, Zoe Weiss, Shannon Kim, Abdullah Bannan, Joseph A. Piccirilli, Jack W. Szostak

## Abstract

Coded ribosomal peptide synthesis could not have evolved unless its sequence and amino acid specific aminoacylated tRNA substrates already existed. We therefore wondered whether aminoacylated RNAs might have served some primordial function prior to their role in protein synthesis. Here we show that specific RNA sequences can be nonenzymatically aminoacylated and ligated to produce amino acid-bridged stem-loop RNAs. We used deep sequencing to identify RNAs that undergo highly efficient glycine aminoacylation followed by loop-closing ligation. The crystal structure of one such glycine-bridged RNA hairpin reveals a compact internally stabilized structure with the same eponymous T-loop architecture found in modern tRNA. We demonstrate that the T-loop assisted amino acid bridging of RNA oligonucleotides enables the rapid template-free assembly of a chimeric version of an aminoacyl-RNA synthetase ribozyme. We suggest that the primordial assembly of such chimeric ribozymes would have allowed the greater functionality of amino acids to contribute to enhanced ribozyme catalysis, providing a driving force for the evolution of sequence and amino acid specific aminoacyl-RNA synthetase enzymes prior to their role in protein synthesis.

## Introduction

In all extant life, amino acids are covalently attached to tRNAs by aminoacyl-tRNA synthetase enzymes to generate the substrates for ribosomal translation. Because RNA- and amino acid-specific aminoacylation is necessary for coded protein synthesis, aminoacyl-RNA synthetase ribozymes likely predated the ribosome(*1*–*3*). Many RNA aminoacylating ribozymes have been isolated by laboratory evolution, supporting this possibility(*4*–*7*). However, their evolution in the RNA World presupposes a selective advantage for the synthesis of aminoacylated RNAs. We have previously demonstrated that the enhanced reactivity of aminoacylated RNAs can accelerate template-directed ligation and facilitate the assembly of active ribozymes composed of chimeric oligomers of RNA and amino acids(*8, 9*). However, nonenzymatic RNA aminoacylation is extremely inefficient(*4, 10*–*13*). Without a moderately effective nonenzymatic pathway for RNA aminoacylation, it is difficult to imagine a selection pressure that could drive the evolution of ribozymes that would enhance RNA aminoacylation. Amino acids are thought to have been widely available on the early Earth(*14*–*16*), and multiple prebiotic amino acid activation pathways have been explored(*11, 17*–*21*) but due to the poor nucleophilicity of the 2′,3′-diol of RNA(*22*), activated amino acids either hydrolyze or polymerize into random peptides before RNA aminoacylation can occur(*11, 12, 17, 18, 23*). The inefficient synthesis of aminoacylated RNA is further compounded by the rapid hydrolysis of aminoacyl esters(*24*–*27*), resulting in low steady state levels of aminoacylated RNA even with repeated amino acid activation.

Induced proximity is an effective strategy for enabling aminoacylation to compete with hydrolysis and peptide formation. For example, in a nicked duplex, aminoacylated RNA can be made in 15 % yield via aminoacyl transfer from a 5′-phosphocarboxy anhydride to the 2′,3′-diol(*28*). Nicked hairpin loop architectures can facilitate the same aminoacyl transfer chemistry, the yield of which is dictated by the sequence of the RNA loop(*29*). However, this route to aminoacyl-RNA synthesis relies on preformed 5′-phosphocarboxy anhydrides, which are even more labile than aminoacyl esters(*12, 18*). Beginning instead with preformed 5′-phosphoramidates, where the N-terminus of an amino acid is linked to a 5′-phosphate, the 2′,3′-diol can efficiently capture the amino acid carboxylate under activating conditions to form chimeric, amino acid-bridged hairpin loops. Aminoacyl-RNA can then be retrieved after acid hydrolysis of the phosphoramidate linkage(*30*). How early life could exploit such a multistep synthesis of aminoacylated RNA is not clear.

Here, we demonstrate that certain RNA sequences can be nonenzymatically aminoacylated and then captured by the formation of amino acid-bridged RNA stem-loops in near-quantitative yield. This process of activated amino acid capture occurs via a two-step reaction pathway in which an activated amino acid carboxylate reacts with the 3′-terminus of an acceptor oligonucleotide, and the amine reacts with the 5′-phosphorimidazolide of a downstream capture strand (**Fig. 1A**). The second step leverages the spatial proximity afforded by the RNA secondary structure to accelerate the ligation. We used deep sequencing to identify 3′-overhang sequences that capture activated glycine efficiently. We solved the structure of one such glycine-bridged stem-loop by X-ray crystallography and found that the structural features necessary for loop-closing ligation closely match those found in T-loops, structures that are universally conserved in tRNA and rRNA. We then demonstrated that the T-loop capture of activated glycine can be used to assemble a functional, chimeric aminoacyl-RNA synthetase ribozyme in high yield and without the need for a template. Our work outlines a sequence of reactions that leverages inefficient nonenzymatic RNA aminoacylation to produce an efficient enzymatic reaction. Similar processes may have played an important role in the primordial assembly of chimeric ribozymes, within which bridging amino acids could then have contributed to enhanced catalysis. We suggest that the ability to assemble ribozymes with precisely positioned internal amino acids could have favored the evolution of sequence and amino acid specific aminoacyl-RNA synthetase ribozymes, thus setting the stage for the evolution of coded peptide synthesis.

**Fig. 1.**
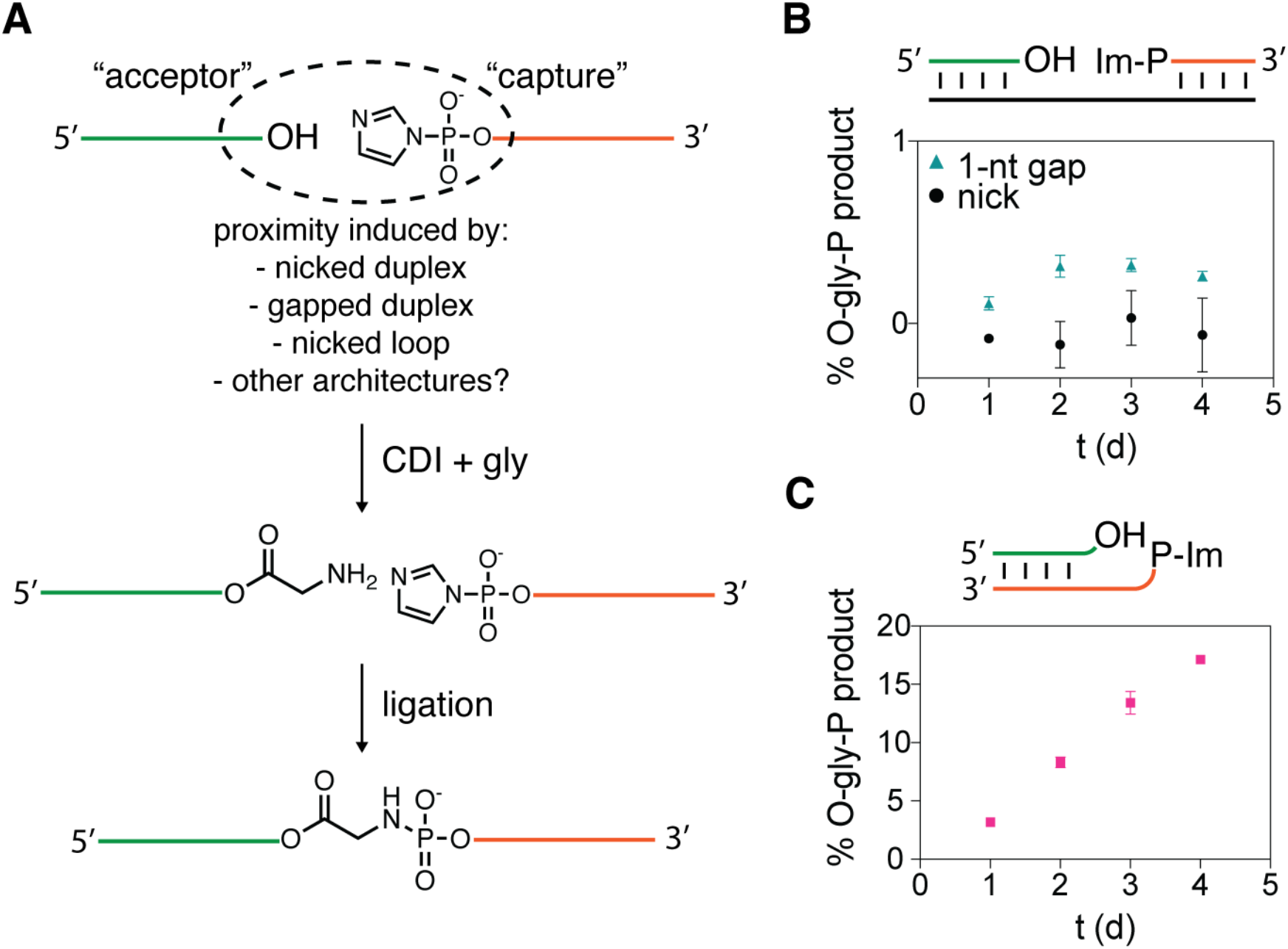
Aminoacylation capture strategy. **A:** General capture scheme that includes an acceptor RNA that is aminoacylated and a capture RNA that is activated as a 5′-phosphorimidazolide (Im-P). **B:** Low yield of aminoacylation capture in a nicked duplex (black circles) or a duplex with a one-nucleotide gap (green triangles). **C:** Aminoacylation capture by a nicked stem-loop construct. All reactions were performed in technical triplicates in 100 mM HEPES pH 8, 5 mM MgCl_2_, 10 μM oligonucleotides, 50 mM premixed CDI + gly added in 12.5 mM increments every 24 h at 0 °C.

## Results

### Proximity-based aminoacyl-RNA capture

Our preliminary experiments confirmed earlier literature results showing very low yields of 2′(3′)-aminoacylated RNA using glycyl-imidazolide prepared from *N,N*′-carbonyldiimidazole (CDI) and glycine (**Figs. S1–2**)(*13, 31*). We have previously shown that a pre-aminoacylated 2′,3′-diol reacts rapidly with a 5′-phosphorimidazolide in a nicked duplex context to yield an amino acid-bridged RNA product(*8*), in which the aminoacyl ester linkage is stabilized by at least two orders of magnitude(*8, 27*). Based on that observation, we expected that the overall yield of nonenzymatic RNA aminoacylation would be increased by capturing *in situ* aminoacylated RNA in an amino acid-bridged RNA product. To our surprise, we were unable to detect any amino acid-bridged product using the same nicked duplex construct when we used CDI-activated glycine to carry out *in situ* aminoacylation (**Figs. 1B, S3**). Operating on the assumption that the poor yield was due to the steric congestion of the nicked RNA substrate, we proceeded to test other potentially more open RNA configurations, such as gapped duplexes, but failed to detect significant product accumulation (**Figs. 1B, S3**).

Serendipitously, while using the Flexizyme ribozyme to aminoacylate the 3′-terminus of a 5′-(2-methylimidazole)-activated oligonucleotide, we noticed the accumulation of an unusually slowly migrating product on acidic urea-PAGE (**Fig. S4A**). Characterization of this product revealed that it was a glycine-bridged hairpin loop, formed by the aminoacylation of one oligonucleotide and ligation to the activated 5′-phosphate of another molecule of the same oligonucleotide (**Fig. S4**). To begin to explore the generality of this type of amino acid-bridged loop-closing ligation, we designed a nicked hairpin loop based on the P2 stem-loop found in the dFx Flexizyme(*6*) and were gratified to observe ligated glycine-bridged product in 17% initial and 30% optimized yield (**Figs. 1C, S5**). To define the activated glycine species responsible for loop-closing ligation, we first tested the *N*-carboxyanhydride of glycine (gly-NCA), a prebiotically accessible intermediate in the CDI-activation scheme (**Scheme S1**)(*32*). Gly-NCA by itself led to no reaction, but loop-closing ligation was restored following the addition of imidazole, suggesting that glycyl-imidazolide is the aminoacylating agent (**Fig. S6**). Failure to form the ligated product after periodate oxidation of the RNA, as well as rapid hydrolysis of the loop-closed product upon alkaline treatment, are consistent with the formation of a glycyl-RNA ester linkage. In addition, high resolution mass spectrometry was consistent with a single glycine ester linkage bridging the diol of the acceptor and the 5′-phosphate of the capture strand (**Fig. S7**).

The stability to hydrolysis of the glycine-bridged loop at pH 8 far exceeded the stability of the same bridge in a duplex context, or a comparable bridge in a single stranded construct(*8*) (**Table S1, Fig. S8**). This result suggested that the structure of the loop could confer increased stability to the labile aminoacyl ester linkage, beyond that conferred by phosphoramidate formation. In addition to glycine, this particular nicked loop architecture captured activated Phe, Ala, and Leu to varying degrees and exhibited moderate stereoselectivity for the L enantiomer, hinting at a potential stereochemical selectivity mechanism based on the RNA architecture (**Fig. S9**).

### Deep sequencing screen for efficient capture sequences

To identify RNA 3′-overhang sequences that efficiently capture activated amino acids, we designed a screen based on high-throughput sequencing (**Figs. 2A, S10**). We first prepared libraries of RNAs with randomized 3′-overhangs of different lengths (4-7 nt), and then subjected the RNA pools to the aminoacylation capture reaction. The purified ligated products were then treated to remove the amino acid (see Methods) and generate an RNA library for suitable for reverse transcription, PCR amplification, and deep sequencing. The resulting distribution of reads, normalized to the distribution in the starting library, suggested that under our reaction conditions all sequences can capture activated glycine at a low background level while a small proportion of sequences confer much more efficient amino acid bridged loop-closing ligation (**Fig. S11**). However, synthesizing and assaying the sequences with the highest number of reads for 4, 5, and 6-nt overhangs did not result in the identification of particularly high-yielding sequences (**Fig. 2B**). In contrast, the 7-nt overhangs with the highest number of reads resulted in a near-quantitative yield of the loop-closed ligation product (**Fig. 2B**).

**Fig. 2.**
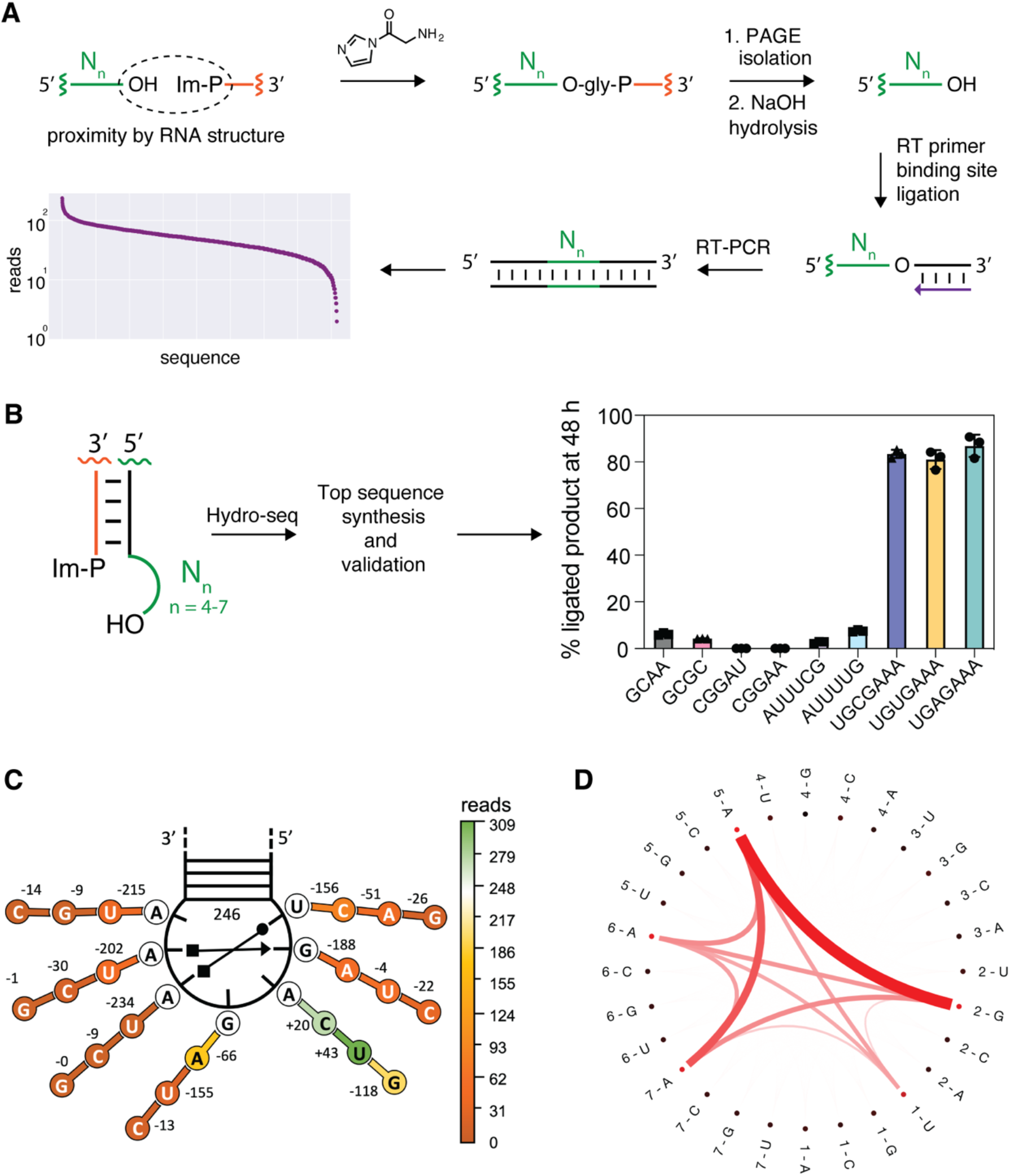
Sequencing assay for the identification of efficient capture sequences. **A:** Schematic of the sequencing assay. A library of acceptor RNA strands consisting of a constant region followed by 4 to 7 randomized nucleotides was incubated with a complementary constant sequence capture strand bearing an imidazole-activated 5′-phosphate, and then chemically aminoacylated with CDI and glycine. Base-pairing of the donor and acceptor strands generates a 5 base-pair stem with a 4 to 7 nucleotide 3′-overhang. Loop-closed products were purified by PAGE and the amino acid bridge selectively hydrolyzed to liberate the acceptor strand. Quantitative enzymatic ligation of the RNA 3′-end to an oligonucleotide that acts as a primer binding site for reverse transcriptase allowed for cDNA synthesis, PCR amplification and deep sequencing. **B:** The capture reaction using the two or three sequences with the greatest number of reads from each stem-loop construct (4–7-nt overhangs on a double-stranded stem). The reaction was performed in technical triplicates in 100 mM imidazole pH 8, 5 mM MgCl_2_, 1 μM oligonucleotides, and 50 mM CDI + gly at 0 °C. **C:** Substitution analysis based on the sequencing data. The change in the number of reads is shown for each nucleotide mutation of the 5′-UGAGAAA-3′ overhang independent of the other mutations. **D:** Median interaction plot as computed from Shapley values in the top cluster. Thickness and color density represent the strength of interaction between pairwise features.

The capture reaction using the 5′-UGAGAAA-3′ overhang reached ≥80% yield after 24 hours even with only 1 mM glycine and CDI, which is 50-fold less than used in all earlier experiments, but failed to produce any product with several other amino acids (**Fig. S12A**,**B**). Interestingly, this overhang sequence produced minimal RNA-only ligation product in the absence of glycine, even with an increased Mg^2+^ concentration and the addition of 1-methylimidazole (**Fig. S12C**), conditions known to afford up to 30 percent ligated RNA product(*33*) in the context of different overhang sequences. These results suggest that there may be distinct steric requirements for different activated amino acids, as well as for the aminoacylation-dependent and independent loop-closing ligation.

To gain insight into the sequence requirements for efficient loop-closing ligation, we examined our entire sequencing data set more closely. The sequencing data for all four overhangs revealed a continuous enrichment in AU-content as a function of reads (**Fig. S13**). An AU-rich sequence composition is known to be associated with higher flexibility and may facilitate bringing the overhang 3′-terminus close to the 5′-phosphorimidazolide of the capture strand(*34, 35*). All single mutations of the 5′-UGAGAAA-3′ overhang at positions 1, 2, 5, 6, and 7 corresponded to sequences that had a lower number of reads (**Fig. 2C**), suggesting that a highly sequence-specific architecture is needed to maximize capture efficiency. We searched for additional sequences that might yield efficient capture by training a booster tree regression model and obtaining estimated impacts of sequence features on the final read counts by employing SHAP analysis. Clustering the Shapley values using a K-means algorithm and synthesizing the second and third best cluster representatives yielded much poorer glycine capture (**Fig. S14**), implying that sequences like 5′-UGAGAAA-3′ possess unique features necessary for efficient capture. To shed light on those features and the synergistic effects in the top cluster sequences, we built a median interaction matrix using the SHAP python package (**Fig. 2D**). Strong positive interactions between 1-U, 2-G, and 5-6-7-A define this top sequence cluster, strengthening the notion that this sequence forms a coordinated architecture that facilitates aminoacylation and ligation.

### Structural elements required for aminoacylation-dependent ligation

In an effort to understand the efficient glycine aminoacylation and loop-closing ligation mediated by the 5′-UGAGAAA-3′ sequence, we set out to determine the structure of the loop-closed product. To accomplish this, we designed an RNA construct that consists of two loops connected by a five base-pair duplex region (**Figs. 3A, S15A**). One loop contains a sequence (5′-AAACA-3′) that binds to the antibody fragment Fab BL3-6(*36*), and the other loop contains the aminoacylated overhang sequence (5′-UGAGAAA-3′). Following aminoacylation and loop-closing ligation, the dumbbell shaped product RNA was used to screen for optimal crystallization conditions. Crystals that diffracted well were obtained, and the structure of the RNA-Fab complex was solved to 2.07 Å resolution with an R_work_ of 22% and an R_free_ of 25% (PDB accession code 9AUS, **Table S2**). The asymmetric unit contained two Fab-RNA complexes, one of which was more ordered than the other and is the subject of the analysis described below.

**Fig. 3.**
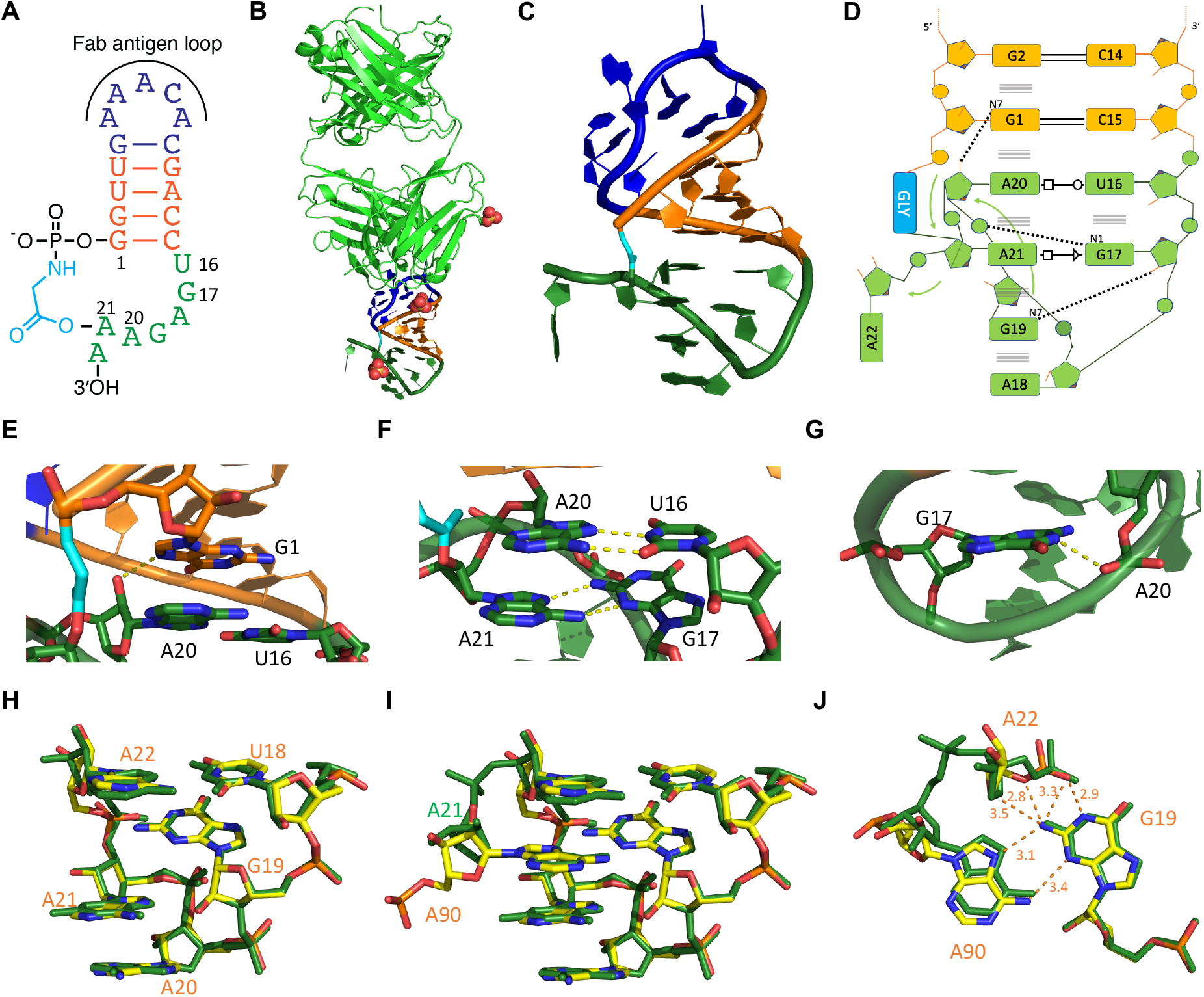
Secondary and tertiary structures of the ligated loop construct determined by x-ray crystallography. **A:** Schematic secondary structure of the overall chimeric dumbbell RNA crystallization construct. **B:** Overall arrangement of the ligated loop construct and Fab BL3-6 in the crystal structure, with the ligated loop in green, glycine linker in cyan, sulphate ions as spheres. **C:** Tertiary structure of the ligated loop construct. **D:** Secondary structure of the ligated loop; interactions are denoted using the Leontis-Westhof nomenclature. **E:** A20:2′OH interacts with G1:N7. **F:** Parallel base pairing interactions between A20 and U16 & A21 and G17 in the ligated loop. **G:** G17:N1 interacts with A20:OP2. **H:** Superposition of ligated loop structural motif (green) and the T-loop motif (yellow) from the FMN riboswitch (PDB 3F2T) without the intercalating base. **I:** as in H but with the intercalating base. **J:** as in I, showing distances and possible interactions between G19, A22 and A90 of the FMN riboswitch.

Interpreting the electron density map was challenging at first, because the distance between the 5′-phosphate of the RNA and the 3′-hydroxyl was far too great to be consistent with the expected glycine bridge. To our surprise, iterative rounds of model building and refinement revealed positive density corresponding to the glycine linker between the 5′-end of the P1 stem (G1) and the 2′-OH of the penultimate residue of the 5′-UGAGAAA-3′ sequence (i.e. A21 of the crystallization construct) (**Fig. S16A**). As a result, the 3′-terminal A22 of the overhang sequence protrudes from the structure of the closed dumbbell-shaped ligated RNA (**Fig. 3C**). This structure was highly unexpected, because of the demonstrated participation of the 2′,3′-diol in aminoacylation for a different, previously studied construct (**Fig. S7**). The 5′-P to 2′-hydroxyl ligated product is a novel RNA branching reaction, mediated by glycine, which leaves the 3′-terminus available as a handle for subsequent binding events and reactions. We carried out several tests to confirm that the site of aminoacylation in the 5′-UGAGAAA-3′ sequence was indeed an internal 2′-hydroxyl and not the expected terminal 2′,3′-cis-diol of the RNA. First, oxidation of the cis-diol by periodate treatment had no effect on the aminoacylation-mediated loop-closing reaction (**Fig. 4**). In contrast, 2′-O-methylation of the penultimate A in the acceptor strand loop significantly inhibited the reaction. In addition, truncation of the overhang to 5′-UGAGAA-3′ had the same effect as 2′-O-methylation of the penultimate A, indicating that only the complete 7-nt overhang can readily adopt the conformation necessary for efficient aminoacylation-dependent loop closing (**Figs. 3–4**).

**Fig. 4.**
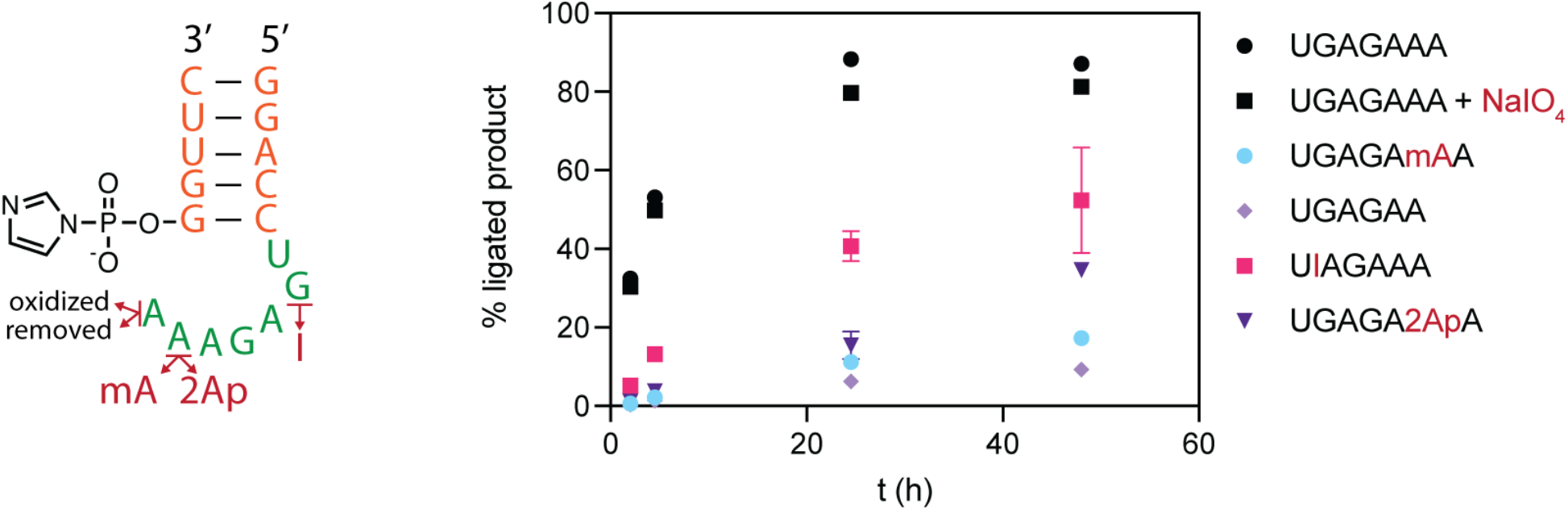
Effect of modifications of the overhang on the loop-closing reaction. The capture reaction was performed with the acceptor RNA modified as indicated. Loop-closing ligation reactions were performed in technical triplicates in 100 mM imidazole pH 8, 5 mM MgCl_2_, 1 μM oligonucleotides, and 50 mM CDI + gly at 0 °C. mA is 2′-OMe-A; I is inosine; 2Ap is 2-aminopurine.

In the crystal structure of the glycine-bridged stem loop, the 2′-aminoacylated loop adopts an architecture that is stacked on and continues the central double-stranded region of the dumbbell shaped RNA. The folded overhang sequence 5′-UGAGAAA-3′ exhibits a number of striking features. The first two nucleotides of the 7-nt overhang, U16 and G17, form consecutive noncanonical base-pairing interactions with the Hoogsteen edges of downstream adenosine residues A20 and A21, continuing the P1 duplex stack and creating a stem-loop-like structure (**Figs. 3C–F**). The string of stacked purines continues below A20 and A21 with G19 and A18, but these nucleotides are not stacked in consecutive order. Rather, the backbone linking G19 and A20 is fully stretched out, leaving a pocket in between these two residues. The backbone then makes an abrupt turn at A20, reversing direction and allowing A21 to be sandwiched into this pocket (**Fig. 3C**). The four nucleotides G17, A18, G19 and then the non-consecutive A21 form the loop that caps the extended base-paired stem (**Fig. 3C,D**). The overall structure of the folded and ligated overhang sequence exhibits a dense network of stacking and pairing interactions, supplemented by three additional hydrogen bonds that likely stabilize the observed configuration (**Fig. 3E–G**). These interactions were evident in our deep sequencing data analysis (**Fig. 2D**), suggesting that future aminoacylation efforts with RNA architectures other than hairpin loops could rely on our computational pipeline to inform on potential RNA structures. Interestingly, residue A18, which makes no nucleobase-specific hydrogen-bonding interactions with other residues (but is stacked on G19), can be replaced with U or C with little effect on the yield of loop-closing ligation (**Fig. 2B–D**).

Since the 5′-UGAGAmAA-3′ construct only partially inhibited the overall product formation, we scaled up the reaction and purified the loop-closed A21 2′-OMe product (**Fig. S15B**). We then crystallized the RNA-Fab complex and determined its structure (PDB accession code 9AUR, **Table S2, Figs. S16B, S17**). In this structure the glycine linker lies between the terminal 5′-phosphate (G1) and the 3′-terminal A22 ribose (**Fig. S17A–D**). However, we cannot resolve whether the aminoacyl group esterifies the 2′-OH or the 3′-OH, or possibly a mixture of the two. Overall, the loop-closed 2′-OMe-modified overhang adopts a structure that is highly similar to that of the unmodified overhang sequence, except that the aminoacylated residue A22 now occupies the pocket previously occupied by the aminoacylated A21 (**Fig. S17E–J**). Conversely, the non-aminoacylated residue A21 protrudes from the folded structure into the solvent, as did the non-aminoacylated residue A22 of the original construct. The tertiary interactions stabilizing the closed-loop structure are the same in both structures, except for the backbone path flanking the aminoacylated residue (**Fig. S17D–J**).

Overall, the backbone and nucleobase arrangements of the first five nucleotides of the overhang sequence (5′-UGAGA-3′) in both structures bear a strong resemblance to the T-loop architecture (**Figs. 3H, S18A**), a ubiquitous motif that frequently mediates tertiary interactions within noncoding RNAs, including the T-loop of tRNA(*37*–*40*). A subclass of these compact U-turn-like structures create a pocket in which a nucleobase can intercalate between nucleobases 4 and 5, forming a continuous base-stack, and can also interact with nucleobase 2 through hydrogen bonding (**Figs. 3I–J, S18B–C**)(*41, 42*). Indeed, within structured RNAs, T-loops can serve as receptors for adenosine residues forming a trans sugar edge/Hoogsteen interaction with a guanosine at position 2 (**Figs. 3I,J**)(*43, 44*), analogous to the A21 or A22 insertions observed in our constructs.

Considering that nicked duplexes and other overhang sequences ligate much less efficiently than the nicked hairpin loop construct, the T-loop architecture observed in our structures may be responsible for facilitating ligation. Alternatively, the folded structure of the ligation product may form after ligation, and not itself be responsible for the enhanced ligation yield. To test the importance for capture efficiency of the sheared pair involving the intercalated A, we prepared two constructs, one in which G17 was replaced with inosine, and one in which A21 was replaced by 2-aminopurine. Both constructs ligated much less efficiently than the original sequence, supporting the hypothesis that the observed tertiary interactions contribute to ligation efficiency (**Fig. 4**).

### Ribozyme assembly by iterated loop-closing ligation

The high yield of the glycine-mediated loop-closing reaction suggested a potential pathway for the structure-guided assembly of ribozymes from small RNA fragments(*45*–*48*). We have previously shown that functional chimeric ribozymes can be assembled via multiple template-directed ligations, but that strategy relies on pre-synthesized aminoacylated RNAs and some means of removing the splint templates so that the ribozyme sequence can fold(*9*). More recently, in a proof-of-principle study, we have shown that an RNA-only loop-closing ligation can be used to assemble hammerhead and ligase ribozymes from two oligonucleotide components (*33*). To investigate whether the glycine-mediated loop-closing ligation could be used to assemble ribozymes without an external template, we divided the dFx Flexizyme into three segments, and placed 5′-UGAGAAA-3′ overhangs at each of the two duplex stems (**Fig. 5A**) (*6, 49*). Upon incubating the modified construct with 50 mM CDI-activated glycine, we observed rapid ligation that yielded the full-length chimeric ribozyme in 43 % yield (**Fig. 5B**). The aminoacylation activity of the isolated chimeric Flexizyme was about half that of the all-RNA ribozyme without amino acid bridges, but 18-fold higher than that of the non-covalently assembled ribozyme (**Fig. 5C**). The high activity of the chimeric Flexizyme suggests that the glycine bridges are well tolerated in the two T-loops, in contrast to their inhibitory effect when placed near the active site(*9*). The efficient aminoacylation-driven untemplated assembly of the Flexizyme points to a possible role for this type of process in the primordial assembly of ribozymes within primitive protocells in which RNA replication is still limited to short unstructured oligonucleotides, and larger structured ribozymes cannot easily be replicated.

**Fig. 5.**
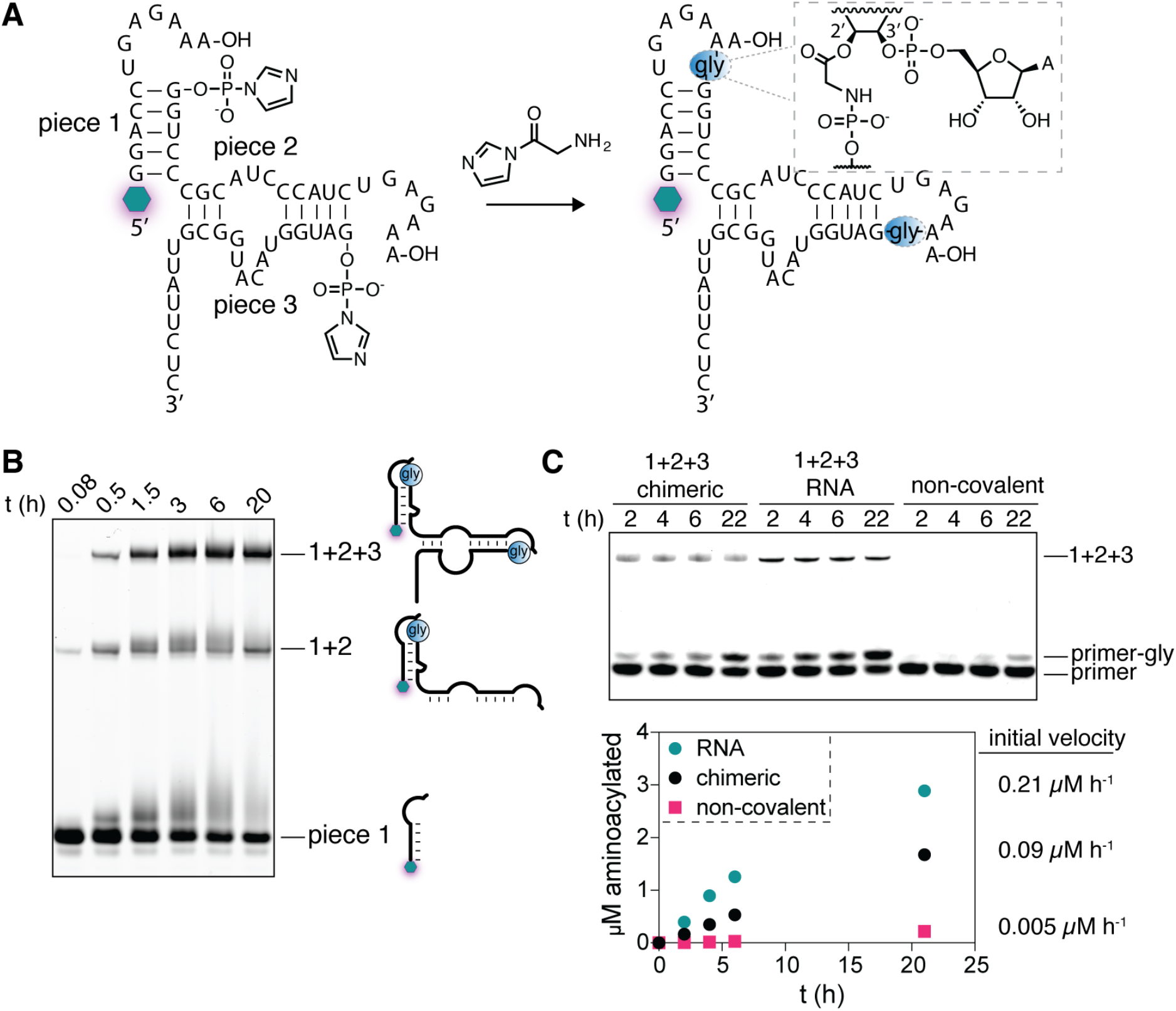
Template-free assembly of a chimeric Flexizyme by two loop-mediated activated glycine capture reactions. **A:** Schematic and secondary structure of the modified Flexizyme ribozyme. The glowing hexagon represents 5′-fluorescein. The blue circles represent the glycine bridge, connecting the RNA as shown in the inset. **B:** Denaturing PAGE of the activated glycine capture reaction over time. The reaction was performed in technical triplicates in 100 mM imidazole pH 8, 5 mM MgCl_2_, 5 μM each oligonucleotide, 50 mM CDI + gly at 0 °C. **C:** Top: denaturing acidic PAGE of the aminoacylation reaction. Bottom: timecourse of the aminoacylation reaction quantified by measuring the ratio of the glycylated vs. not glycylated gel band in each lane. The reaction was performed in technical triplicates in 100 mM imidazole pH 8, 5 mM MgCl_2_, 5 μM primer oligonucleotide, 0.5 μM ribozyme, 5 mM DBE-gly (20 % overall DMSO) at 0 °C.

## Discussion

Our work addresses the possibility that RNA aminoacylation served a primordial function that was advantageous at an early stage of the RNA World, which could help to explain the evolutionary origins of the translation system. We have previously shown that RNA aminoacylation can lead to enhanced rates of template-directed RNA ligation, but achieving enhanced ligation yields required pre-aminoacylated RNAs, which we had to generate by ribozyme-catalyzed aminoacylation. The low steady state levels of RNA aminoacylation that can be achieved by all known pathways for nonenzymatic aminoacylation have made it difficult to imagine how a biologically advantageous function could result from purely chemical aminoacylation. We have now found that specific RNA 3′-overhang sequences can be efficiently aminoacylated in the presence of amino acids that are activated as imidazolides. Attack of the amine of the amino acid on the activated 5′-phosphate of the complementary strand of the duplex, combined with aminoacyl ester formation, results in loop closing ligation and stabilization of the otherwise labile aminoacyl ester. Thus, the presence of both carboxyl and phosphate activating chemistry can lead to the formation of covalently closed hairpin loops with a bridging amino acid at the ligation junction. We have shown that this template-independent mode of ligation can be iterated to generate larger structured RNAs with ribozyme activity, including a ribozyme that is itself an aminoacyl-RNA synthetase. We suggest that similar processes may have acted within protocells at an early stage of the RNA World to enhance the assembly of ribozymes from sets of short oligonucleotides, such as those that might constitute a fragmented genome(*50*).

While the mechanism by which the 5′-UGAGAAA-3′ sequence facilitates the aminoacylation and loop-closing ligation reaction remains unclear, several arguments suggest that the unligated overhang sequence may be pre-organized in a manner similar to the product structure, and that this structure catalyzes one or both reactions. First, the site of aminoacylation was, to our surprise, the internal 2′-hydroxyl of the penultimate (-1 position) nucleotide of the overhang, and not the expected cis-diol of the 3′-terminal nucleotide. The -1 nucleotide is intercalated between two nucleotides of the T-loop structure, possibly positioning its 2′-hydroxyl in a manner favoring its aminoacylation and/or facilitating subsequent attack of the glycine amine on the nearby activated 5′-phosphate. In addition, the loop-closing ligation reaction is highly specific for glycine. Finally, destabilization of the folded structure by chemical mutagenesis greatly reduces the yield of loop-closing ligation. Thus, preorganization of the 3′-overhang sequence in the folded geometry provides potential explanations for both the site and amino acid specificity of the loop-closing reaction. The crystal structure of the ligated product also provides a potential explanation for the increased stability of the aminoacyl ester linkage, because the folded structure restricts the solvent accessible surface area of the ester by 90%, which would not occur in duplex or unstructured strands (**Fig. S19**).

The chimeric glycine-bridged RNA folds into a stable compact structure with the same 3-D architecture as seen in the canonical T-loop that is widely distributed in biological RNA structures, including the eponymous T-loop of tRNAs. This motif is so small and simple that it is likely to have evolved independently many times wherever it was needed to fulfill a structural or, as our results now suggest, a catalytic role. This raises the question of whether other, perhaps more complex and less widely distributed, structural motifs might also exist, some of which might also facilitate aminoacylation and ligation reactions. We are currently screening new RNA libraries for internal loop and bulge-closing ligation reactions, using the same type of deep sequencing assay that allowed us to screen 3′-overhangs for enhanced loop-closing ligation. The identification of sequences that enable additional structure-generating ligation reactions would greatly increase the diversity of RNA structures that could be assembled as a result of aminoacylation and ligation. We are particularly interested in the identification of RNA sequences that confer different amino acid specificities on the aminoacylation reaction, as this could allow for the strategic positioning of bridging amino acids within a folded RNA such that the amino acid side chains could stabilize or modulate the folded structure, or even enhance the catalytic abilities of an assembled ribozyme. While the self-assembly of such chimeric RNAs involves intrinsic sequence and amino acid specificity, these constraints might be reduced if the aminoacylation step was carried out by distinct ribozymes. We therefore suggest that the assembly of chimeric ribozymes with enhanced activities could result in a selective pressure for the evolution of ribozyme aminoacyl-RNA synthetases that are both sequence and amino acid specific, thus generating the substrates later coopted for the evolution of coded peptide synthesis.

## Acknowledgements

Our crystallographic work is based upon research conducted at the Northeastern Collaborative Access Team beamlines, which are funded by the National Institutes of Health grant P30 GM124165. This research used resources of the Advanced Photon Source, a U.S. Department of Energy (DOE) Office of Science User Facility operated for the DOE Office of Science by Argonne National Laboratory under Contract No. DE-AC02-06CH11357. The authors are grateful to Daniel Krochmal for technical support, and Dr. Daniel Duzdevich, Dr. Benjamin Colville, Dr. Long-fei Wu, and Dr. Victor Lelyveld for help with setting up the deep sequencing screen. The authors also thank Caroline Kaminsky, Dr. Saurja Dasgupta, Dr. Jian Zhang, Dr. Filip Boskovic, Dr. Aman Agrawal, Prof. Yamuna Krishnan, and the Szostak laboratory for their thoughtful suggestions on the manuscript.

## Author contributions

Conceptualization: AR, JWS

Methodology: AR, AL, MT, HRMA, ZW, SK, AB, JAP, JWS

Investigation: AR, AL

Visualization: AR, AL, MT, HRMA, ZW

Funding acquisition: JAP, JWS

Project administration: AR, AL, JWS, JAP

Supervision: AR, AL, JWS, JAP

Writing: AR, AL, JWS, JAP

## Data and materials availability

The BL3-6 Fab expression vector is available upon request. PDB accession codes for the loop-closed glycine-bridged dumbbell RNA and the A21 2′-OMe modified dumbbell RNA are 9AUS and 9AUR, respectively. The code for general deep sequencing analysis can be found at: https://github.com/szostaklab/aminoacylation/blob/main/Hydro-seq_example.ipynb. The remaining data are available in the main text or the supplementary materials.

## Supplementary Materials

Materials and Methods

Scheme S1

Figs. S1 to S19

Tables S1 to S3

References (*51-63*)

