## Supplementary Information for "Structure-guided aminoacylation and assembly of chimeric RNAs"

**The PDF file includes:**

Materials and Methods
Scheme S1
Figs. S1 to S19
Tables S1 to S3

### Materials and Methods

General information.

All reagents were purchased from Sigma-Aldrich and Fisher Scientific, unless otherwise noted. Oligonucleotide synthesis reagents were purchased from ChemGenes and Glen Research. Illumina sequencing reagents were purchased from Illumina and New England Biolabs.

Common protocols and recipes.

Ethanol precipitation: 0.1 volumes of 3 M sodium acetate pH 5.5 and at least 3 volumes of ethanol were added to the RNA solution. The precipitation was allowed to proceed for 20 minutes on dry ice before being centrifuged at maximum speed at 4 ºC. The supernatant was removed, and the pellet was washed twice with cold 80 % ethanol.

Isopropanol precipitation: 0.1 volumes of 5 M ammonium acetate and at least 3 volumes of isopropanol were added to the RNA solution. The precipitation was allowed to proceed for 20 minutes on dry ice before being centrifuged at maximum speed at 4 ºC. The supernatant was removed, and the pellet was washed twice with cold 80 % ethanol.

Soaking buffer: 5 mM sodium acetate pH 5.5 mixed with 2 mM EDTA pH 8.0. The measured pH of this solution was 7 without adjustments.

Acidic soaking buffer: 5 mM sodium acetate pH 5.5 mixed with 2 mM EDTA pH 8.0. The pH was adjusted to 5.0 with 1 M HCl.

Quenching buffer: 50 mM EDTA pH 8.0, 1x bromophenol blue, 95 % v/v formamide.

Acidic quenching buffer: 10 mM EDTA pH 8.0, 1x bromophenol blue, 100 mM sodium acetate pH 5.0, 150 mM HCl, 75 % v/v formamide.

Oligonucleotide synthesis.

All oligonucleotides were synthesized on the K&A H-6 solid-phase DNA/RNA synthesizer. Randomized nucleotides in the sequencing constructs were introduced from a hand-mixed solution containing 3:3:2:2 A:C:G:U phosphoramidites. Following cleavage and deprotection as recommended by the manufacturer, the oligonucleotides were evaporated in a speed-vac for 2 hours and lyophilized for 16 hours. Removal of the 2′-O-TBDMS groups was performed in 100 µL DMSO and 125 µL triethylamine trihydrofluoride at 65 ºC for 2.5 hours. Oligonucleotides were then precipitated with isopropanol, dissolved in 5 mM EDTA in 99 % v/v formamide, and purified by denaturing urea-PAGE. The desired gel bands were cut, crushed up, and soaked in soaking buffer for 16 hours. Finally, the oligonucleotides were filtered and desalted with either C18 Sep-Pak cartridges (Waters) for lengths $<25$ nt or Amicon MWCO filters for lengths $>25$ nt. Analysis of pure RNA was performed on an Agilent 6540 mass spectrometer.

Oligonucleotide activation.

To a solution of 250 µM RNA in 125 mM imidazole pH 7 was added EDC.HCl, so that the final 300 µL solution contained 200 µM RNA, 100 mM imidazole pH 7, and 100 mM EDC.HCl. The reaction was rotated for 2 hours at 22 ºC before being precipitated with 30 µL acetone saturated with sodium perchlorate and 1 mL acetone on dry ice for 10 minutes. After centrifugation and supernatant removal, the pellet was washed twice with a 1:1 v/v mixture of acetone and diethyl ether. Activation efficiency was determined by HPLC using the Atlantis TM T3 column (3 µm, 4.6 x 150 mm) at a flow rate of 0.5 mL/min. The following gradient was used: (A) aqueous 50 mM triethylammonium acetate (pH 7.0) and (B) acetonitrile, from 6% to 12% B over 20 min. Activation efficiency was routinely over 85 %.
Activation with 2-methylimidazole was performed as described previously(*8*).

Model monomer aminoacylation (Figure S1).
Model MeO-pA was synthesized following the reported procedure of Ivanovskaya and coworkers(*51*). Aminoacylation was attempted using either (i) CDI + glycine or (ii) gly-NCA(*52*) + imidazole, in D2O at pD = 7.5 (imidazole buffer). Experiments were incubated and monitored using time course 1H NMR, sampling every 10 min for 4 h.
A: To a solution of glycine (94 mg, 1.25 M, 50 eq.) in D2O (1 mL) was added carbonyldiimidazole (244 mg, 1.5 M, 60 eq.) and the mixture was rapidly stirred for 2 min. Next, 100 µL of the aforementioned solution was transferred to a solution of MeO-pA (6.25 mM) in imidazole buffer (pD = 7.5, 625 mM in D2O, 400 µL). The combined mixture was vortexed for 30 s before being transferred to an NMR tube. The progress of the reaction was monitored by 1H NMR spectroscopy over the course of 4 h.
B: To a solution of MeO-pA (6.25 mM) in imidazole buffer (pD = 7.5, 625 mM in D2O, 400 µL) was slowly added a solution of gly-NCA (1.25 M in d 6 -DMSO, 100 µL, 50 eq.) with vigorous stirring. The combined mixture was vortexed for 1 min before being transferred to an NMR tube. The progress of the reaction was monitored by 1H NMR spectroscopy over the course of 4 h.
C: An authentic sample of 2′/3′-glycyl product for 1H NMR comparison was prepared following the procedure of Sutherland and coworkers(*29*).

Model oligonucleotide aminoacylation (Figure S2).
100 µL of a 0.5 M glycine solution in water were added to 9.75 mg (1.2 eq) of CDI and rapidly vortexed for 20 seconds. This solution was then added to a solution of RNA such that the final 50 µL reaction contained 5 µM RNA, 360 mM buffer, 40 mM $MgCl_2$, and 100 mM premixed gly + CDI. 1 µl aliquots were taken at the indicated time points, quenched with 9 µL acidic quench buffer and analyzed by acidic 20 % denaturing urea-PAGE. Acidic PAGE was made with 100 mM sodium acetate pH 5.0 instead of the usual 1x Tris-Borate-EDTA, and it was run in this same buffer at 25 W for 2.5 hours at 4 ºC. The remaining reaction was precipitated with ethanol, desalted with a zip-tip C18 column, and injected into the Agilent 6540 mass spectrometer.

Flexizyme-catalyzed oligonucleotide aminoacylation (Figures S2, S4, and 5C).
To a solution of RNA and Flexizyme dFx(*6*) or the modified dFx_S7 (Table S3) in imidazole pH 8.0 and $MgCl_2$ was added dinitrobenzyl ester of glycine (DBE-gly) so that the final

concentrations were 5 μM RNA, 0.5 μM Flexizyme, 100 mM imidazole pH 8.0, 5 mM $MgCl_2$, and 5 mM DBE-gly (20 vol% DMSO). Aminoacylation efficiency was analyzed as above.

Amino acid capture reaction (Figures 1, S3, S5-9, S12, S14).

The following standard conditions were used unless otherwise noted: to a solution of 5.55 mM $MgCl_2$, 111 mM imidazole pH 8.0, 11.1 μM acceptor oligonucleotide, 11.1 μM capture oligonucleotide in 18 μL was added 500 mM of premixed gly + CDI in 0.5 μL portions every 24 hours over 96 hours at 0 ºC. The reaction was quenched every 24 hours by mixing 0.5 μL aliquots with 4.5 μL of quenching buffer and analyzed by 20 % denaturing urea-PAGE. The percent product was obtained by quantifying the per-lane normalized band intensity in the ImageQuant TL software.

The NaOH treatment shown in Figure S7B was performed by treating 4 μL of the quenched 96 hour timepoint with 1 μL of 1 M NaOH for 3 minutes at 22 ºC. The hydrolysis was quenched by the addition of 1 μL of 1 M HCl.

For the LC-TOF analysis in Figure S7C, a 25x scale reaction run for 96 hours was concentrated with Amicon MWCO filters, precipitated with ethanol, desalted with a zip-tip C18 column, and injected into the Agilent 6540 mass spectrometer.

Hydrolysis reaction (Table S1, Figure S8).

Synthesis: 500 μL reaction containing 5 mM $MgCl_2$, 100 mM imidazole pH 8.0, 10 μM acceptor oligonucleotide, 10 μM capture oligonucleotide, and 50 mM premixed gly + CDI was incubated for 96 hours at 0 ºC. The reaction was concentrated with Amicon MWCO filters to 50 μL, diluted with 50 μL formamide, and purified by 20 % denaturing urea-PAGE at 4 ºC. The desired product band was isolated, crushed, and soaked in acidic soaking buffer for 3 hours at 4 ºC. The solution was then filtered, concentrated with Amicon MWCO filters to 50 μL, and precipitated with ethanol.

Hydrolysis: 10 μL reactions containing 2.5 μM of the amino acid-bridged construct for the loop-closing reaction and an additional 50 μM duplexing oligonucleotide (A2C2duplex or A3C3duplex in Table S3) for the nicked duplex reaction were heated at 90 ºC for 1 minute followed by slow cooling (0.1 ºC/s) to 22 ºC. The annealed reactions were then diluted with HEPES pH 8.0 and $MgCl_2$ to give the final conditions in 10 μL: 200 mM HEPES pH 8, 2.5 mM $MgCl_2$, 0.75 μM annealed construct. At the indicated time points, 1 μL aliquot was diluted with 14 μL quenching buffer containing 2.4 μM of the reverse complement of the duplexing oligonucleotide (A2C2revcomp or A3C3revcomp in Table S3), heated at 95 ºC for 3 minutes, quickly cooled on ice, and analyzed by 16 % denaturing urea-PAGE. The ratio of the full-length glycine-bridged construct to the hydrolyzed product was obtained by quantifying the per-lane normalized band intensities in ImageQuant TL. Negative natural logarithm of the ratio of the remaining bridged construct (P) to the initial bridged construct ($P_0$; assumed to be 1) at each time point was plotted against time. The slope represented the $k_{obs}$, and the half-lives were obtained by dividing ln(2) with the $k_{obs}$.

Deep sequencing (Figures 2, S10, S11).

Preparative reaction:

1. To a solution of 7.5 mM $MgCl_2$, 150 mM imidazole pH 8.0, 1.5 µM each acceptor oligonucleotide construct, 1.5 µM capture oligonucleotide per acceptor construct, 1.5 µM protecting oligonucleotide per construct in 2.666 mL was added 150 mM of premixed gly + CDI in 334 µL portions at 0 and 24 hours at 0 ºC. At 48 hours, the reaction was concentrated using Amicon MWCO filters to 50 µL, diluted with 50 µL formamide, and purified by 16 % denaturing urea-PAGE at 4 ºC. The product band was crushed, soaked in acidic soaking buffer for 3 hours, filtered, concentrated with Amicon MWCO filters, and desalted using the Zymo Oligo Clean and Concentrator kit.
2. The isolated product was then hydrolyzed in 200 mM NaOH for 3 minutes, quenched with HCl, diluted with formamide, purified by 16 % denaturing urea-PAGE, and isolated as above.
3. The hydrolyzed product was then ligated to the preadenylated RT primer binding oligonucleotide using the standard NEB protocol for T4 RNA ligase 2, truncated KQ with 10 % PEG at 25 ºC for 22 hours. To a 20 µL ligation reaction was added 0.5 µL Proteinase K and the reaction was incubated for 15 minutes at 22 ºC. The reaction was cleaned up with 2 washes of 25:24:1 v/v/v phenol:chloroform:isoamyl alcohol followed by desalting with the Zymo Oligo Clean and Concentrator kit.

At this step, the initial library that did not undergo steps 1 and 2 was also subjected to steps 3-7. This was done to capture the initial library sequence bias and correct for it by performing a normalization of the sequences that captured glycine.

4. The ligated product was reverse transcribed using the NEB ProtoScript® II First Strand cDNA Synthesis Kit and the cDNA was isolated using the Zymo Oligo Clean and Concentrator kit with the recommended alkaline RNA removal.
5. The cDNA (200 ng) was then PCR amplified and multiplexed with the NEB Q5 polymerase kit and NEBNext® Multiplex Oligos for Illumina® (Index Primers Set 1) using the standard Q5 PCR conditions for 6 cycles.
6. The amplified product was purified by 1.4 % agarose gel and extracted with the Monarch® PCR & DNA Cleanup Kit.
7. Prior to pooling, the final concentration of each sample was determined with the 4200 Agilent Tapestation. The samples were then pooled to the final total concentration of 1 nM and prepared for NGS per the Illumina protocol for Miseq, with a 30 % PhiX spike-in.

At steps 1, 2, and 3, 1 µL aliquots were diluted with quenching buffer and analyzed by 16 % denaturing urea-PAGE as in Figure S10.

Analytical reactions with individual sequences: the reactions were performed exactly as in step 1 above, except the reaction was scaled down 100x. Instead of continuing with the sequencing workflow, 1 µL aliquots were quenched at 48 hours in 9 µL quenching buffer and analyzed by 16 % denaturing urea-PAGE. The percent product was obtained by quantifying the per-lane normalized band intensity in the ImageQuant TL software.

General deep sequencing data analysis (Figures 2, S11).

The detailed and annotated custom python code for each construct analysis can be found at the following github link: https://github.com/szostaklab/aminoacylation/blob/main/Hydro-seq_example.ipynb.

The raw reads were first filtered for quality, the length of the randomized region, and the homemade barcode where applicable. The reads were then trimmed to only display the randomized regions of interest. The reads of the glycine capture reaction were then normalized as follows: the overall number of reads for the initial library was divided by the number of unique sequences observed to obtain the optimal read per sequence if the library were truly random; reads for each unique sequence in the initial library were then divided by the optimal read per sequence number to obtain the normalization factor; finally, reads for each unique sequence in the glycine capture dataset were divided by the normalization factor to obtain the normalized read count for each unique sequence. Sequences with less than 5 reads in the initial library were omitted from the analysis and assigned the read count of 0. This normalization procedure removes biases that occurred during the initial random library synthesis. The sequences and their corresponding read counts were then plotted.

<u>In-depth deep sequencing data analysis (Figures 2, S13-14).</u>

Most of the entries in our datasets correspond to poorly represented sequences that we interpret as weakly reacting. For example, the read counts in our sequencing dataset for heptamers had a mean equal to 20, corresponding to a mean ligation yield of ~ 4%. We are particularly interested in the sequences with the high number of reads and on the features that differentiate these sequences from the bulk.

To dissect the impact of the sequence content on the reads, we studied the 7 nt-long loop results. The original sequencing dataset for 7 nt-long loops included 15765 sequences out of the 16384 possible heptamers formed by the canonical 4 nucleobases of RNA. The missing sequences were treated as zero-reads sequences, so that they are reintroduced in our expanded dataset to be taken into account for downstream analysis.

To extract quantitative features from the deep sequencing of the 7-nt loops, we trained a boosted tree regression model using Python XGBoost package(*53*). The dataset was split in training/test sets (80/20 split) and the sequences were One-Hot-Encoded to feed the model, so that every one of our 7-nt sequences resulted in a 7 x 4 matrix that was further flattened into a 28 elements-long array. The XGBoost model was trained with {max depth: 6, learning_rate: 0.1, subsample: 0.5, tree_method: 'hist'} and using our test set for early stopping (200 early stopping rounds) to reduce overfitting. Final RMSE for training set and test set were respectively equal to 7.9 and 12.5. The Shapley values were extracted from the trained model using Python SHAP package(*54*).

Before performing clustering on the Shapley values, we decided to modify the dataset by setting every Shapley value for zero-categorical features as equal to 0. Since most of the Shapley value variance belongs to categorical features equal to one (90% of Shapley values for zero-categorical features are between -2.8 and 1.3, while 90% of Shapley values for one-categorical values are between -3.9 and 8.3), we retain most of the model's predictive power. Moreover, since zero-categorical features on average lower the read values predicted from the SHAP analysis, this effect can be partially compensated by assigning a net penalty of ~10 to our reads. By discarding

the zero-categorical features Shapley values, we ended up trading some performance of the model, with overall RMSE increasing from 10.0 to 14.9 (without net subtraction) or 11.1 (with net subtraction), while allowing for better interpretable clusters.
Clustering was performed using the K-means algorithm implemented in scikit-learn(*55*), with the optimal number of clusters chosen using the elbow method(*56*) as equal to 8.
To prepare data for visualization and ease the visualization of clusters with widely uneven size, we subsampled the modified Shapley dataset to contain no more than 300 entries per cluster. To generate the low-dimensional visualization of this reduced Shapley values dataset, we applied the UMAP algorithm implemented in Python UMAP package with neighbor number set to 150. Pairwise interactions between sequence features were extracted using the SHAP package, and the resulting interaction network was visualized as a schemaball with a modified version of Oleg Komarov MATLAB code(*57*).

Amino acid capture with optimized UGAGAAA overhang sequence (Figures 3, 4, S12).
The loop-closing amino acid capture reaction, incubated at 0 ºC, contained the following components in 20 µL: 5 mM $MgCl_2$, 100 mM imidazole pH 8.0, 1 µM acceptor oligonucleotide, 1 µM capture oligonucleotide, and a range of concentrations between 50 mM and 1 mM of premixed amino acid and CDI.

Periodate oxidation (Figures 4, S7).
10 µM of acceptor oligonucleotide was treated with 10 mM sodium periodate in 200 µL for 1 hour at 0 ºC. The oligonucleotide was desalted using Amicon MWCO filters. The 50 µL solution was precipitated with ethanol.

Synthesis of the amino acid bridged constructs for X-ray crystallography (Figure S15).
Dumbbell RNA (Figure 3): 80 µM acceptor A56, 80 µM capture C9, 25 mM EDTA, 100 mM imidazole pH 8, and 12.5 mM of premixed gly + CDI were mixed to a final volume of 8 mL and incubated for 48 hours at 0 ºC. The reaction was then concentrated using Amicon MWCO filters to 150 µL, diluted with 150 µL formamide, and purified using 20 % denaturing urea-PAGE. The desired amino acid-bridged product gel band was crushed, soaked in acidic soaking buffer for 3 hours at 4 ºC, filtered, desalted with C18 Sep-Pak cartridges, and lyophilized. The lyophilized product was then circularized to complete the Fab-binding loop using the NEB T4 RNA Ligase 1 (ssRNA Ligase), High Concentration under standard conditions for 60 minutes at the final reaction volume of 30 mL. The ligation reaction was stopped with 240 µL of Proteinase K for 15 minutes. The RNA solution was extracted twice with 25:24:1 v/v/v phenol:chloroform:isoamyl alcohol followed by desalting with the C18 Sep-Pak cartridges. After lyophilization, the RNA was dissolved in 200 µL water and further desalted by precipitation with isopropanol.
The A21 2′-OMe dumbbell construct (Figure S16): 80 µM acceptor A57, 25 mM EDTA, 100 mM imidazole pH 8, and 12.5 mM of premixed gly + CDI were mixed to a final volume of 8 mL and incubated for 48 hours at 0 ºC. The reaction was then concentrated using Amicon MWCO filters to 150 µL, diluted with 150 µL formamide, and purified using 20 % denaturing urea-PAGE. The desired amino acid-bridged product gel band was crushed, soaked in acidic soaking buffer for 3 hours at 4 ºC, filtered, desalted with C18 Sep-Pak cartridges, and lyophilized.

Note: the circularized, amino acid-bridged product migrates faster than the corresponding linear amino acid-bridged RNA on the 20 % denaturing urea-PAGE.

Fab Purification:
The BL3-6 Fab expression vector (available upon request) was transformed into 55244 chemically competent cells (www.atcc.org) and grown on LB plates supplemented with carbenicillin at 100 μg mL-1. Nine colonies from the plate were chosen and inoculated to a starter culture with 100 μg mL-1 carbenicillin, which was grown at 30 °C for 8 hours. Once the starter culture reached an OD 600, 15mLs of starter culture was used to inoculate 1L of 2×YT media and grown for 24 h at 30 °C. The cells were then pelleted via centrifugation at RT, and the cell pellet was resuspended in 1L of freshly prepared CRAP-Pi media supplemented with 100 ug mL-1 carbenicillin. The cells were set to grow for 24 h at 30 °C, harvested via centrifugation at 4 °C and frozen at -20 °C. Frozen cell pellets were lysed in PBS supplemented with 0.4 mg mL-1 of Lysozyme and 0.01 mg mL-1 of DNase I. After 30 minutes PMSF was added to a final concentration of 0.5 mM. After 30 minutes, the mixture containing cellular debris and lysate was centrifuged, 45 min, 12000 rpm, rotor type JLA 16.250 (Beckman) at 4 °C. Lysate was transferred to new sterile bottles and centrifuged again for 15 minutes, 12000 rpm, at 4 °C. Supernatant was filtered through 0.45 μm filters into a sterile bottle (Millipore Sigma, www.sigmaaldrich.com), and Fab proteins were purified using the AKTAxpress fast protein liquid chromatography (FPLC) purification system (Amersham, www.gelifesciences.com) as described previously(*36, 58*). The lysate in PBS (pH 7.4) was loaded into a protein A column, and the eluted Fab in 0.5 M acetic acid was buffer exchanged back into the PBS (pH 7.4) using 30kDa cutoff Amicon filter and loaded into a protein G column. The Fab was eluted from protein G column in 0.1 M glycine (pH 2.7) and then buffer-exchanged into 50 mM NaOAc, 50 mM NaCl buffer (pH 5.5) and loaded into a heparin column. Finally, the eluted Fab in 50 mM NaOAc, 2 M NaCl (pH 5.5) was dialyzed back into 1× PBS (pH 7.4), concentrated, and analyzed by 12% SDS-PAGE using Coomassie Blue R-250 staining for visualization. Aliquots of Fab samples were tested for RNase activity using the RNaseAlert kit (Ambion, www.thermofisher.com). The aliquots of Fab samples were flash frozen in liquid nitrogen and stored at −80 °C until further use.

Crystallization:
The crystallization construct was analyzed by electrophoretic mobility shift assay under non-denaturing conditions. In the presence of a stoichiometric amount of Fab chaperone, BL3-6 RNA construct shifted quantitatively to the bound state. The same shift did not occur for a non-Fab sample. Validated RNA construct in ultrapure $H_2O$ was subjected to crystallization with no further procedures of refolding. 480 μg of the RNA was mixed with a 1.1 molar equivalent of Fab BL3-6. The complex was then concentrated to 6 mg/mL final concentration of RNA (80 μL). 100 nL + 100 nL hanging drop crystal trials were set in commercially available crystallization kits from Hampton Research and Jena Bioscience using the Mosquito liquid handling robot (SPT Labtech) and allowed to grow for two to three weeks at 4 °C. First crystals grew in 3 days. Loop-closed dumbbell RNA crystallized in Peg Ion: 0.2 M Lithium sulfate monohydrate; 20% w/v Polyethylene glycol 3,350, pH 5.7. A21 2′-OMe dumbbell construct also crystallized in Peg Ion:

0.07 M Citric Acid; 0.03 BIS-TRIS Propane; 16% w/v Polyethylene glycol 3,350, pH 3.8. Crystals were looped and frozen in liquid nitrogen.

Crystallographic data processing (Figures S16, S19):

Diffraction data was collected at APS beam line 24-ID-C. Data sets were integrated and scaled using the on-site RAPD automated programs (https://rapd.nec.aps.anl.gov). The structures were solved using molecular replacement of the Fab BL3-6 from the previously reported structure (PDB code: 7SZU(*59*)) as search model in Phenix Phaser(*60*). Two copies of the RNA and Fab were discovered in the P 1 space group for the loop-closed dumbbell RNA. Only one copy of the A21 2′-OMe dumbbell construct RNA and the Fab was discovered in the unit cell in the P 1 21 1 space group. Using the initial phases from the molecular replacement solution, the RNA was able to be built into the emerging density after multiple rounds of refinement using Coot and Phenix Refine(*61–63*). Visualization and solvent accessible surface area (SASA) evaluation were conducted with PyMOL (The PyMOL Molecular Graphics System, Schrödinger, LLC).

Ribozyme assembly (Figure 5A,B).

The ribozyme assembly reaction, incubated at 22 ºC, contained the following components in 20 µL: 5 mM $MgCl_2$, 100 mM imidazole pH 8.0, 5 µM piece 1, 5 µM piece 2, 5 µM piece 3, and 50 mM premixed gly + CDI. At each time point, 1 µL aliquot was diluted in 9 µL quenching buffer and analyzed by 16 % denaturing urea-PAGE. The percent product was obtained by quantifying the per-lane normalized band intensities in ImageQuant TL.

**Scheme S1**: CDI-mediated glycine activation generates both NCAs and aminoacyl imidazolides. Shown are two possible paths (red and black) to NCA from the aminoacyl urea imidazolide intermediate(*32*).

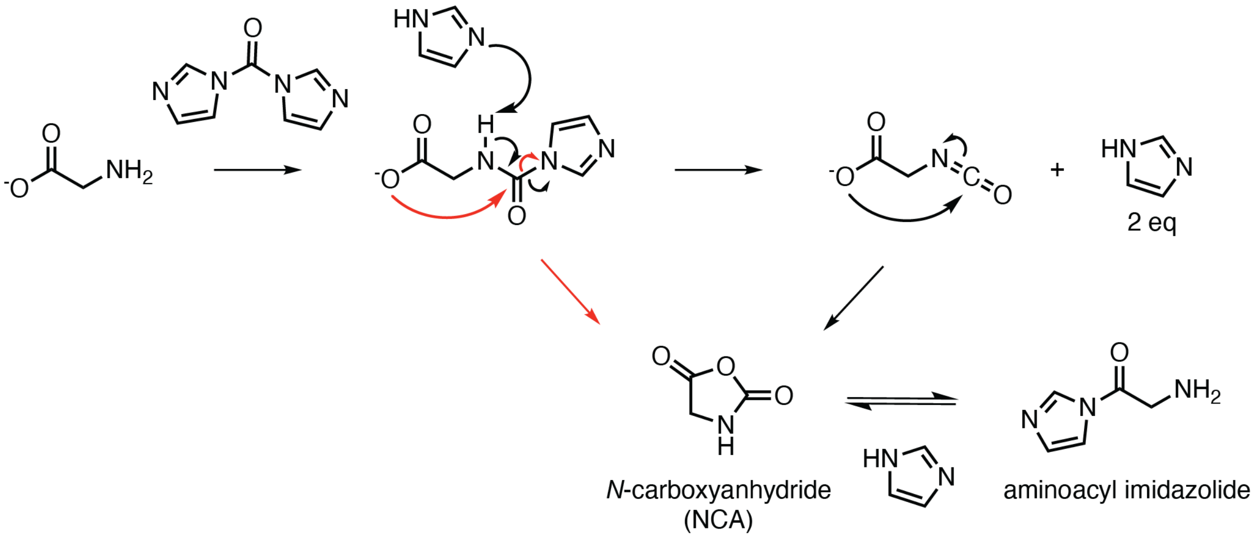

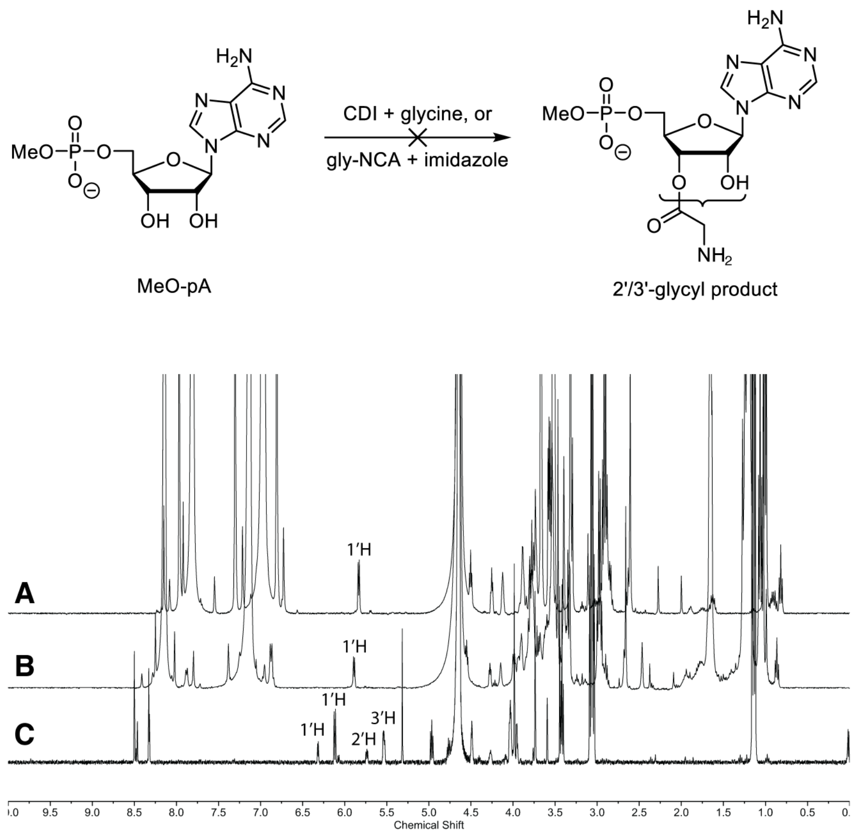


**Fig. S1. Aminoacylation of a model monomer, adenosine 5′-(O-methylphosphate).** Representative $^{1}$H NMR after 2 h of attempted aminoacylation of monomeric nucleotide (MeO-pA). **A:** No aminoacylation was observed in the reaction with premixed CDI + glycine; **B:** No aminoacylation was observed in the reaction with gly-NCA + imidazole; **C:** authentic product (2′/3′-glycyl-5′-MeO-pA). The region of the spectrum between 5 and 6.75 ppm contains the diagnostic signals of successful aminoacylation: 1′H is shifted downfield and appears as two signals due to the 2:1 equilibrium ratio of 3′-glycyl and 2′-glycyl ester. 2′H and 3′H are shifted downfield in the presence of glycyl ester and they also reflect the 2:1 equilibrium.

**A**

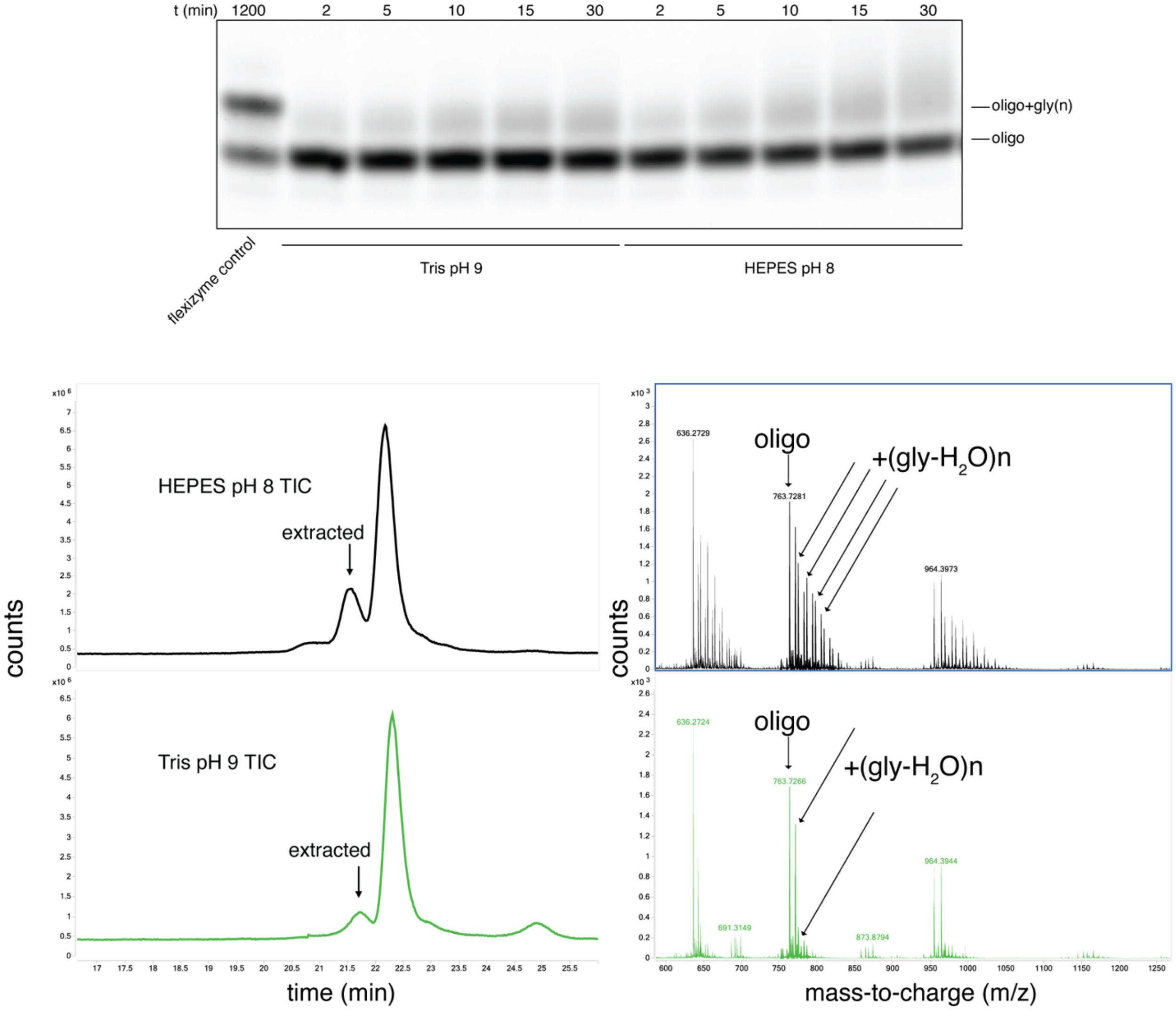


**B**

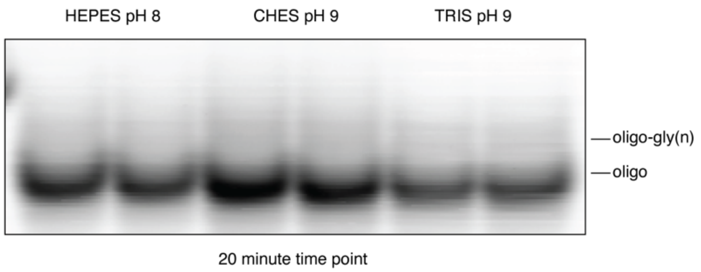


**Fig. S2. Aminoacylation of a model oligonucleotide. A** (top): Acidic PAGE of a reaction timecourse between the model oligonucleotide (FX3 in Table S3) and CDI + glycine added at time point 0. The left-most lane contains a 1200-minute time point of the Flexizyme aminoacylation, which adds a single glycine on the 3′-terminus. **A** (bottom): Analysis of the same reaction by LC-TOF. The newly appearing signal in the total ion chromatogram (TIC) was extracted and exact masses were calculated for the labeled ions in the labeled charge envelope. Top left panel is the TIC of the aminoacylation reaction performed in HEPES pH 8. The top right panel is the extracted ion chromatogram (EIC) of the signal that appeared after the aminoacylation reaction. The exact masses were calculated for the labeled ions, and they corresponded to the non-aminoacylated model oligonucleotide FX3 and the FX3 containing one or more glycyl esters. For clarity, only signals of the ions with the same charge are labeled. The bottom two panels are the same as the top ones, except the aminoacylation was performed in Tris pH 9. Under these conditions, only a singly glycylated FX3 can be observed. **B**: Acidic PAGE of the 20-minute time point of the same reaction as above, except the model oligonucleotide contained all internal deoxyribonucleotides, with only the 3′-terminal ribonucleotide (dFX3 in Table S3).

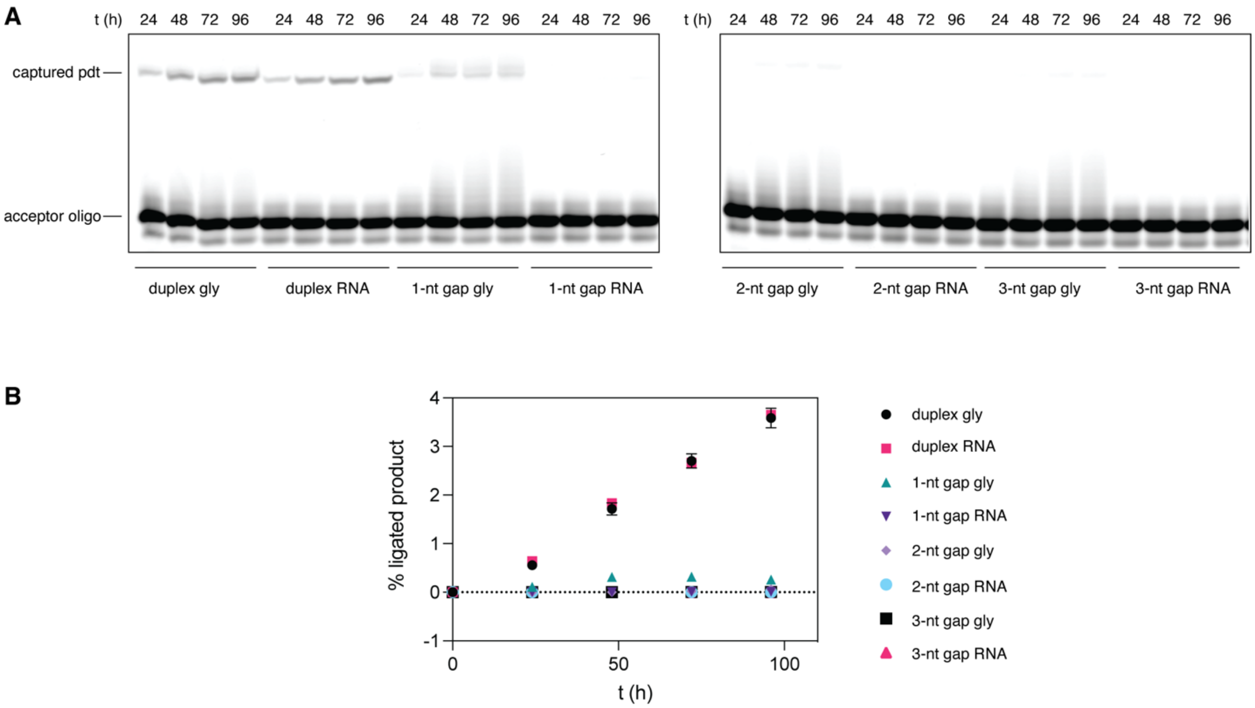


**Fig. S3. Activated amino acid capture by duplexed RNA.** **A**: PAGE analysis of the time-course of the capture reaction with a nicked duplex and 1–3-nt gap-containing duplex. **B**: Quantification of the percent ligated or captured product from the gels in **A**. For each lane, the ratio of the ligated product band (top) to the sum of the ligated product and the starting material acceptor band (bottom) was determined and converted to percent product by multiplying it with 100 %.

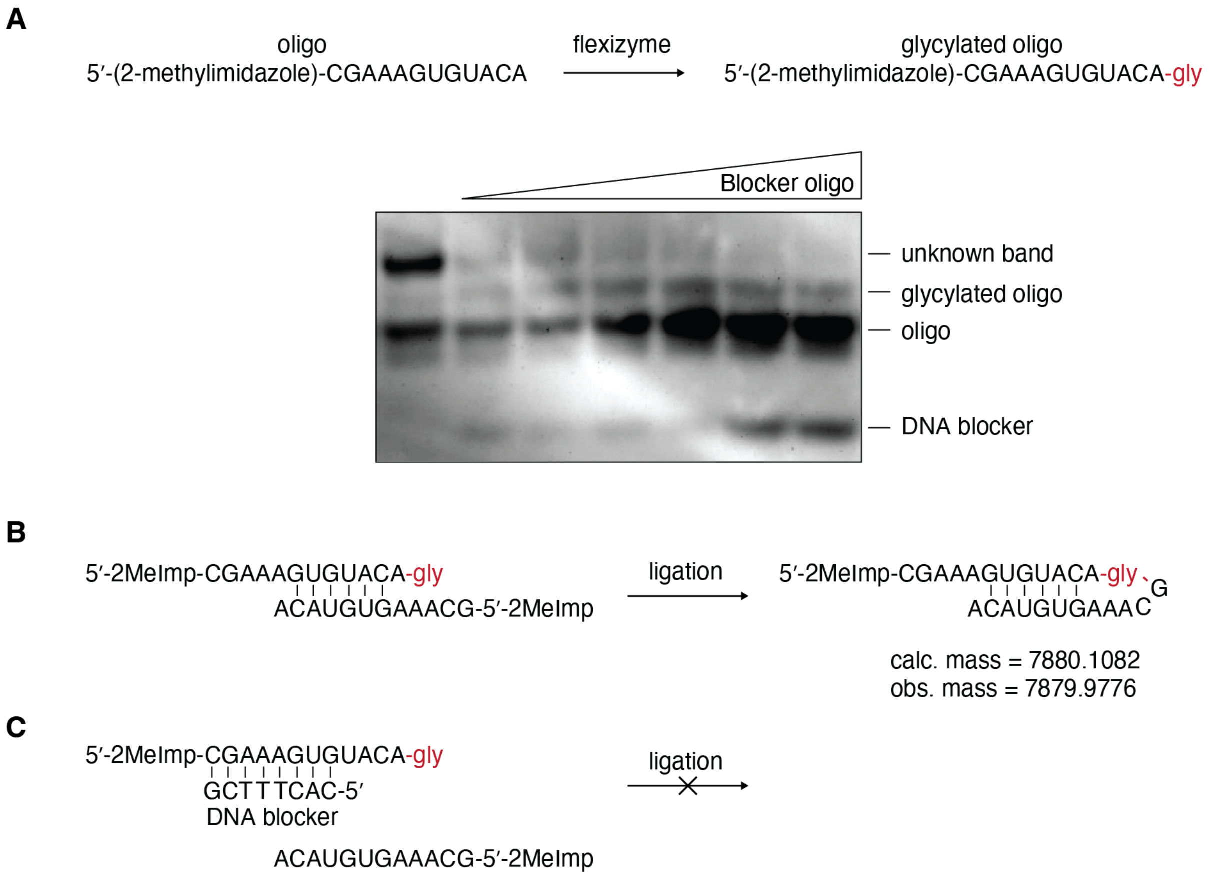


**Fig. S4. Flexizyme-catalyzed aminoacylation of a partially self-complementary oligonucleotide. A**: The general reaction scheme and expected product. Acidic PAGE analysis of the reaction with and without an increasing concentration of a blocker oligonucleotide. **B**: The hypothesized loop amino acid capture due to partial self-complementarity of the activated and aminoacylated oligonucleotide. LC-TOF analysis of the reaction confirming the appearance of the hypothesized product. **C**: Scheme of the blocker oligonucleotide preventing the amino acid-bridged loop formation.

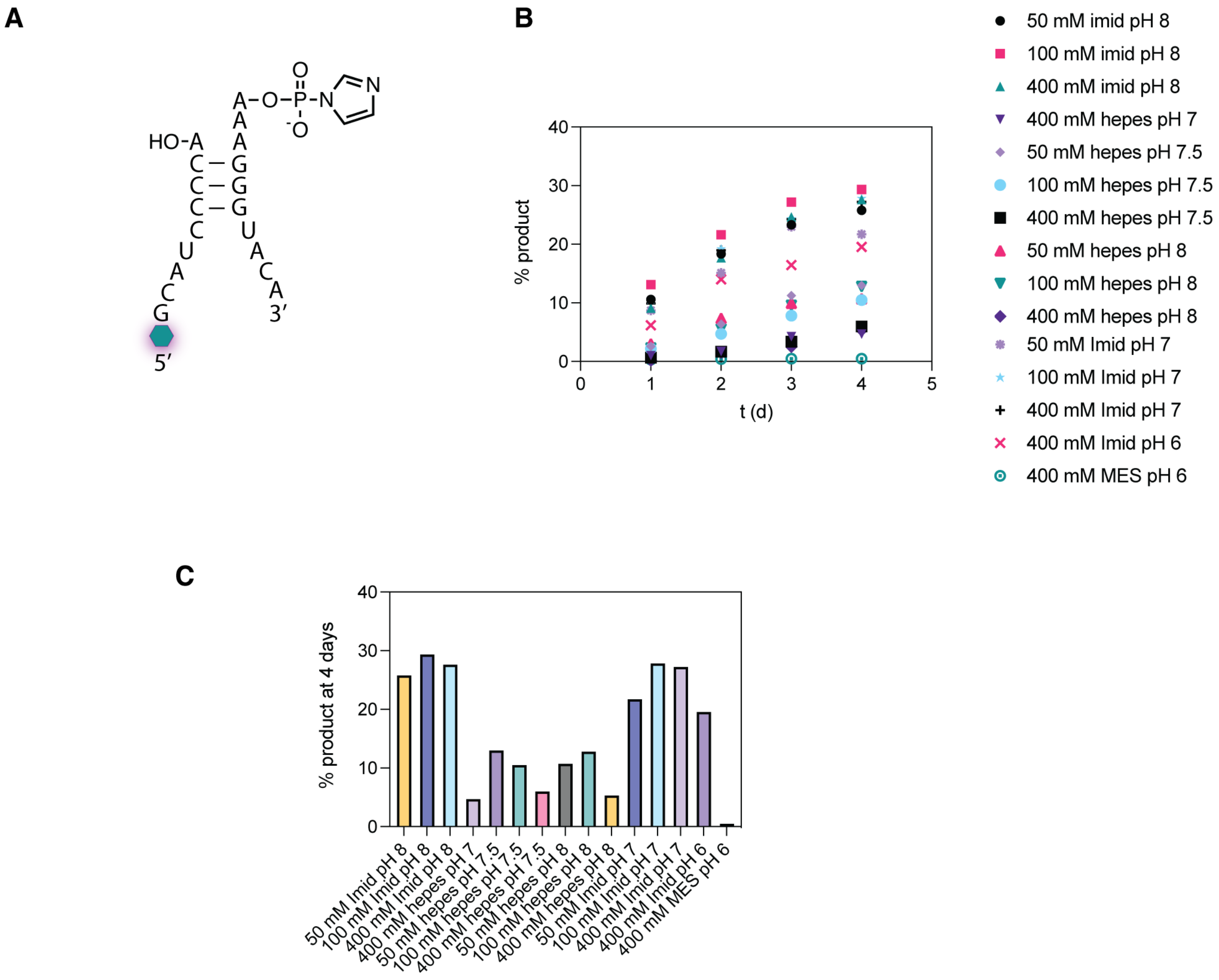


**Fig. S5. Optimization of the pH of the capture reaction using the modified P2 stem-loop of the Flexizyme. A**: Diagram of the nicked stem-loop based on the P2 stem-loop found in the Flexizyme. **B**: Timecourse of the capture reaction at different concentrations of buffer and at different pHs. % product is quantified based on the per-lane normalized band intensity. **C**: Bar graph of the 4-day timepoint from **B** showing the maximum yield at each concentration and pH.

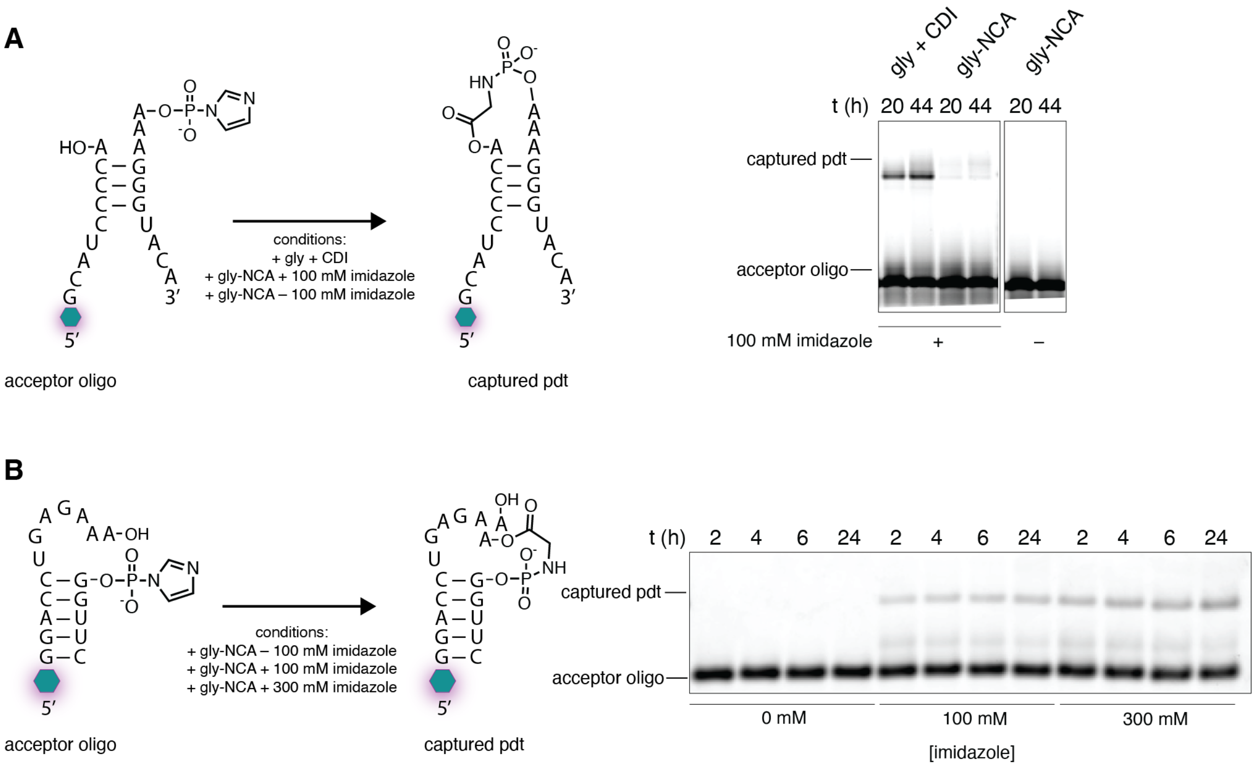


**Fig. S6. Imidazole-catalyzed gly-NCA capture using the modified P2 stem-loop of the Flexizyme and the T-loop containing RNA. A**: On the left is the diagram of the capture reaction. On the right is PAGE analysis of the capture reaction. The conditions for the reactions were: 10 µM each oligonucleotide, 5 mM $MgCl_2$, 100 mM HEPES pH 8, and activated glycine as follows. Gly + CDI = 50 mM premixed gly and CDI; gly-NCA = presynthesized gly-NCA added instead of gly + CDI at 50 mM with or without 100 mM imidazole pH 8. **B**: On the left is the diagram of the capture reaction using the highly efficient T-loop containing RNA that we identified in our sequencing screen. Glycine is bridging the 2′-OH of the penultimate A of the 5′-UGAGAAA-3′ motif and the 5′-phosphate of the downstream G, as shown in Figure 3. On the right is PAGE analysis of the capture reaction. The conditions for the reactions were: 1 µM each oligonucleotide, 5 mM $MgCl_2$, 100 mM HEPES pH 8, and activated glycine as follows. 50 mM of the presynthesized gly-NCA was added to the reaction with the increasing concentration of imidazole pH 8.

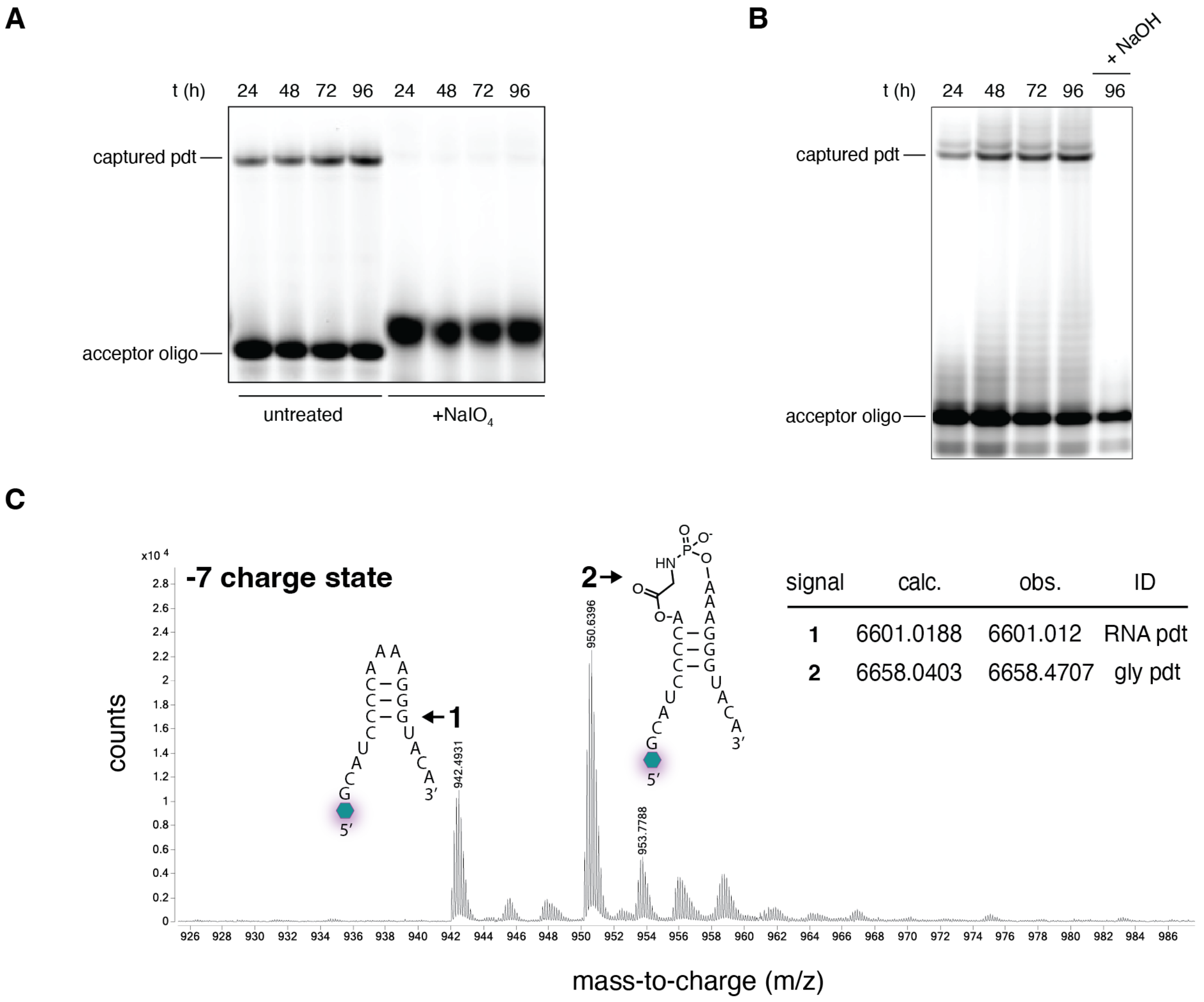


**Fig. S7. Loop-closing amino acid capture reaction characterization using the modified P2 stem-loop of the Flexizyme. A**: PAGE analysis of the capture reaction with and without treatment of the acceptor strand with 10 mM $NaIO_4$. **B**: PAGE analysis of the capture reaction with the 96 h time point untreated and treated with 100 mM NaOH for 2 minutes. **C**: LC-TOF analysis of the crude 96 h reaction. Shown is the extracted ion chromatogram of –7 charge state of the reaction product. Signal 1, "RNA pdt", is consistent with the product of the background RNA only reaction, which is beyond the detection limit of PAGE. Signal 2, "gly pdt", is consistent with the glycine-bridged loop product.

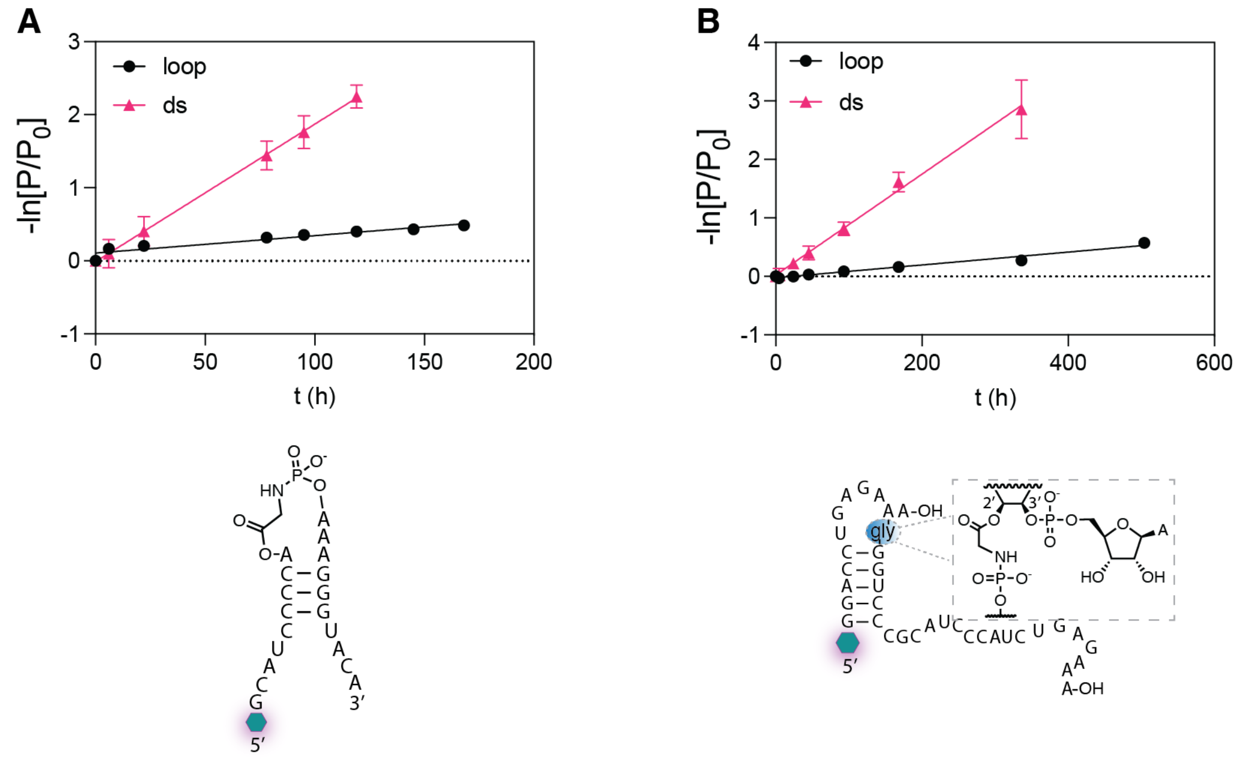


**Fig. S8. Kinetic plots of the glycine-bridged RNA degradation reactions.** Negative natural logarithm of the ratio of the remaining product (P) to the initial product ($P_0$; assumed to be 1) at each time point. **A**: Hydrolysis of the glycine-bridged modified P2 stem-loop of the Flexizyme. Shown below the plot is the diagram of the loop structure. The structure was forced into the double stranded (ds) form by adding the complementary strand in 20-fold excess and performing an annealing cycle. **B**: Hydrolysis of the glycine-bridged modified P1 stem-loop of the Flexizyme containing the 5′-UGAGAAA-3′ T-loop motif. Shown below the plot is the diagram of the loop structure with glycine bridging the 2′-OH of the penultimate A of the 5′-UGAGAAA-3′ motif and the 5′-phosphate of the downstream G, as shown in Figure 3. The double stranded form was formed as above.

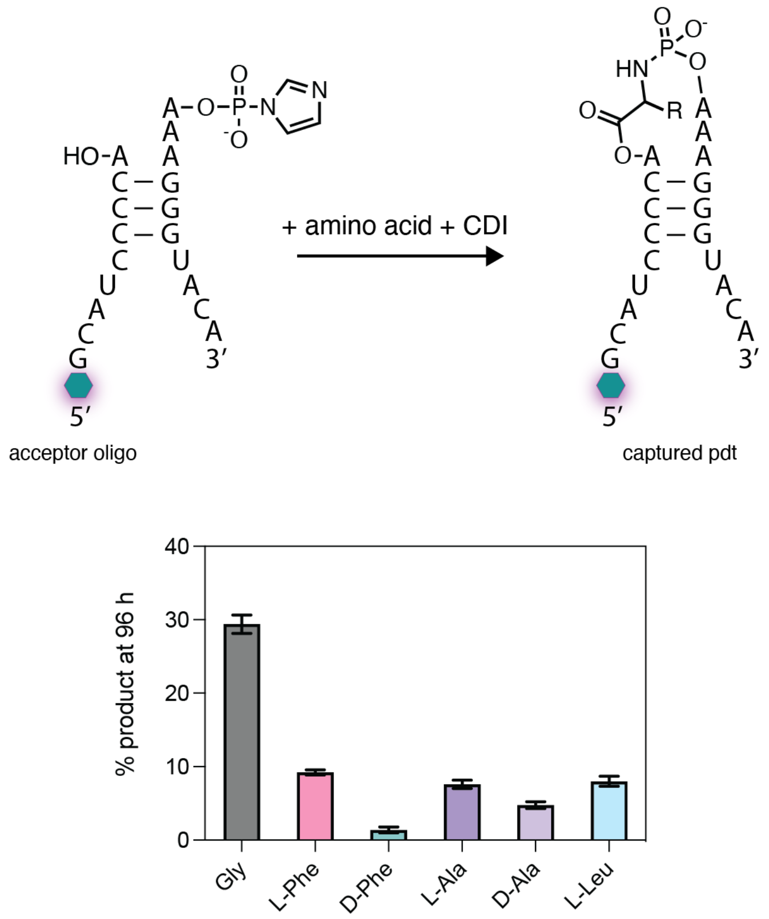


**Fig. S9. The capture reaction across amino acids for the modified P2 stem-loop of the Flexizyme.** Shown is the final yield (at 96 hours) of the capture reaction, quantified by PAGE analysis based on the per-lane normalized band intensity. Standard conditions of 100 mM imidazole pH 8, 5 mM $MgCl_2$, 10 µM each oligonucleotide, 50 mM final concentration of premixed amino acid + CDI (aliquots containing 0.25 µmol of the CDI-activated amino acid were every 24 hours beginning at 0 hours) at 0 ºC were employed.

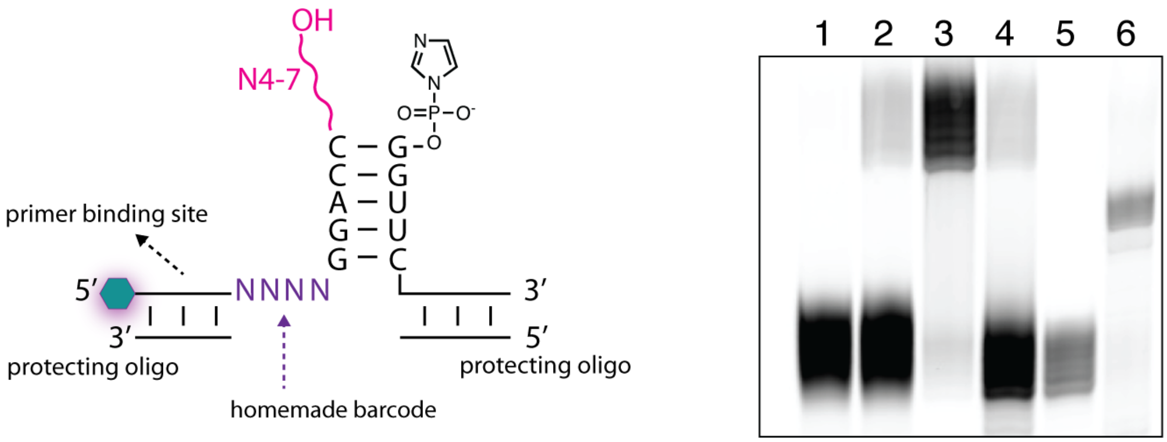


**Fig. S10. Schematic of the sequencing construct and quality controls.** Nicked loop construct, containing a short 5-bp stem and a variable length (4-7 nt) randomized 3′-overhang (see Table S3 for the exact oligonucleotides). The glowing hexagon represents 5′-fluorescein. The encoded primer binding site allowed us to skip the ligation of the 5′-adapter. The protecting oligo was used to prevent any unwanted base-pairing interference. The acceptor oligonucleotide contained a homemade barcode to allow for facile filtering of sequences containing different overhang lengths during data analysis. On the right is a PAGE quality control analysis: lane 1 = reaction without gly showing no RNA-only product; lane 2 = reaction with gly + CDI; lane 3 = the product of the reaction with gly + CDI purified by PAGE; lane 4 = the purified product hydrolyzed with 100 mM NaOH for 3 minutes; lane 5 = PAGE-purified hydrolyzed product; lane 6 = RT primer binding site ligated to the PAGE-purified hydrolyzed product.

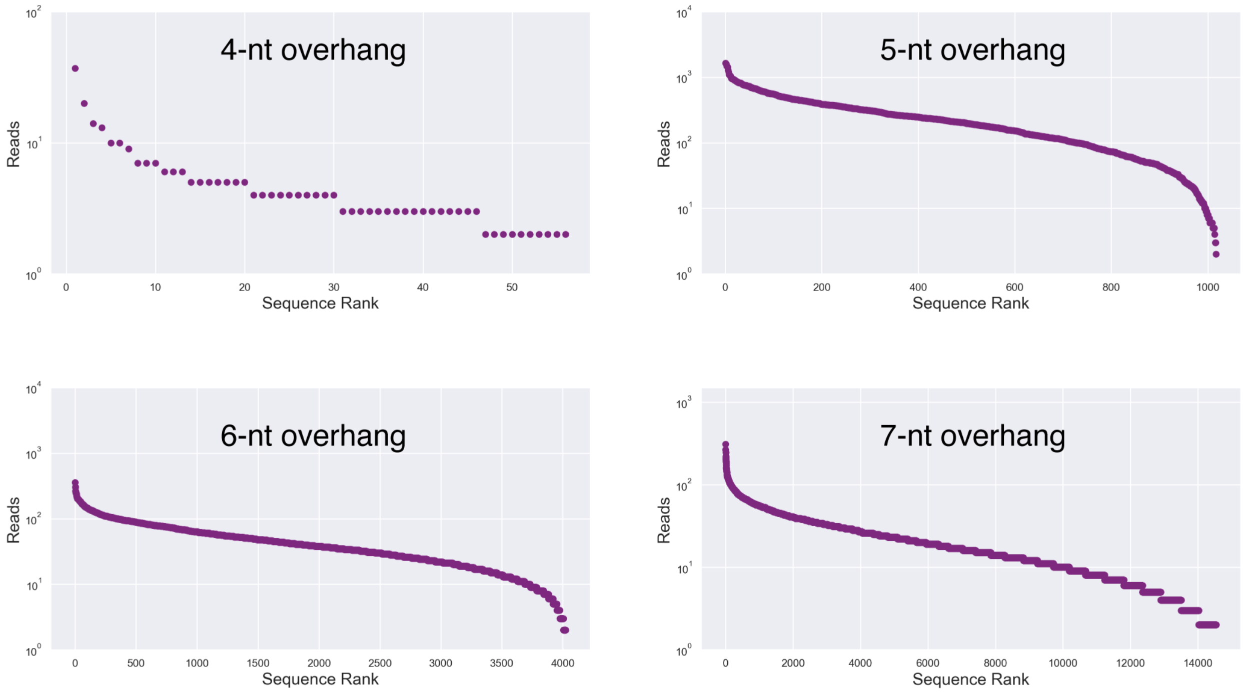


**Fig. S11. The distribution of sequencing reads across sequence space for the nicked stem-loop constructs.** Filtered and sorted sequences plotted as Reads (log-scale) normalized to the distribution of reads in the starting library versus Sequence Rank. Each purple dot represents a unique sequence.

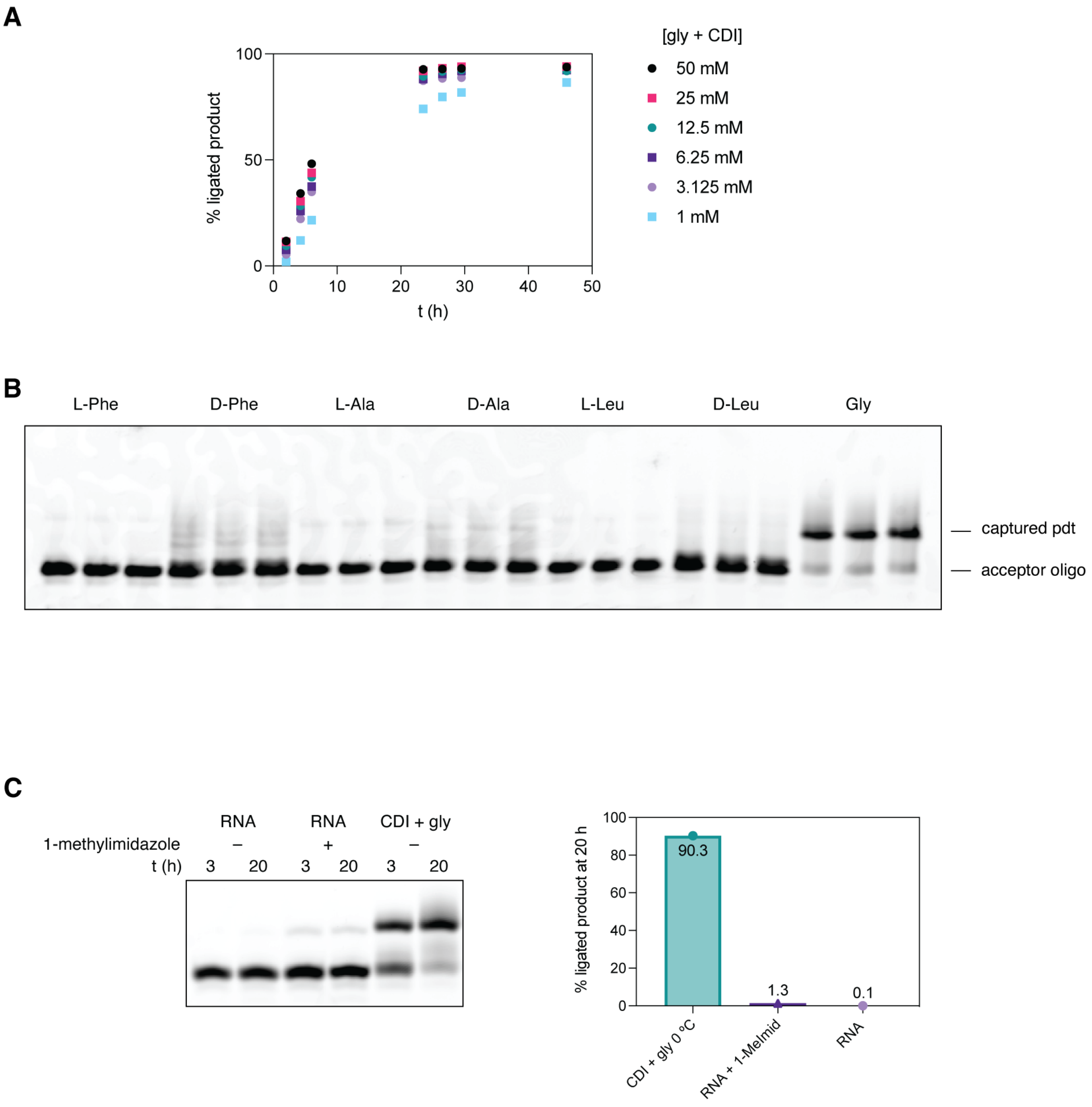


**Fig. S12. Capture of a variety of amino acids and ligation of RNA with the 5′-UGAGAAA-3′ nicked loop. A**: Shown is the capture reaction yield over time with a serial dilution of the premixed gly + CDI under standard conditions. **B**: PAGE analysis of the 96-hour time point of the capture reaction with a variety of activated amino acids. Each amino acid reaction was loaded in triplicates. **C**: RNA-only ligation using the UGAGAAA overhang and optimized RNA-only ligation conditions with and without the addition of the 1-methylimidazole catalyst. RNA-only conditions were 100 mM Na-HEPES pH 8, 50 mM $MgCl_2$, 1 µM each oligonucleotide, and 100 mM optional 1-methylimidazole at 22 ºC. The conditions for amino acid capture were 100 mM imidazole pH 8, 5 mM $MgCl_2$, 1 µM each oligonucleotide, and 50 mM amino acid + CDI at 0 ºC.

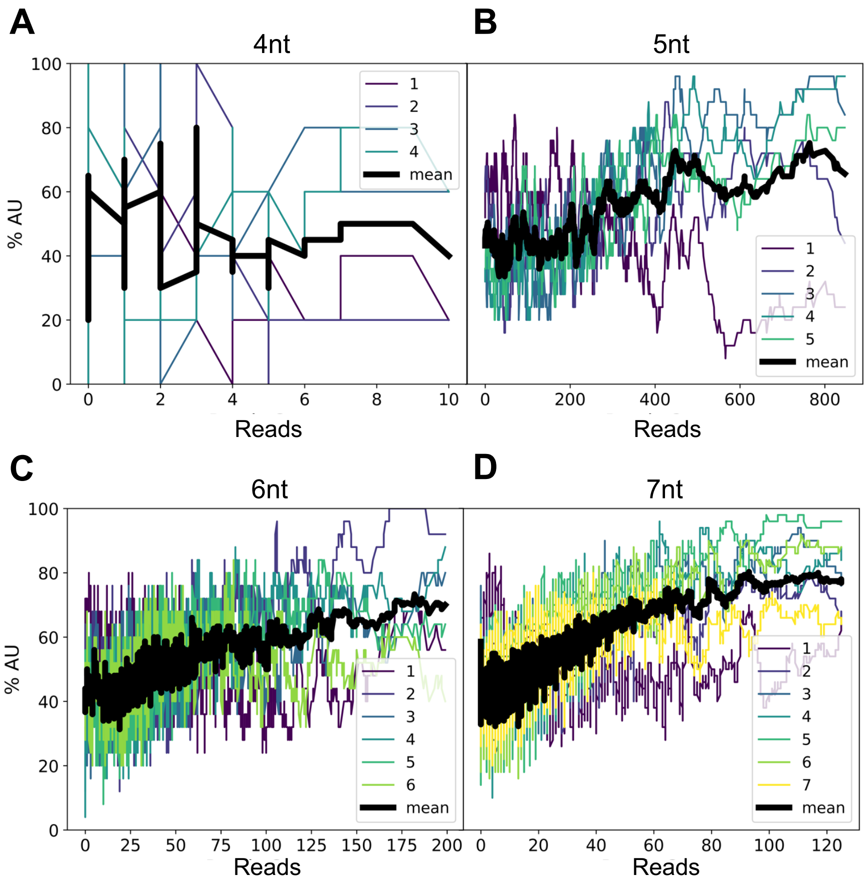


**Fig. S13**. **AU-content as a function of reads for (A) 4-nt long loops, (B) 5-nt long loops, (C) 6-nt long loops, and (D) 7-nt long loops.** Colored traces show values calculated on a per-nucleotide basis, while the black trace shows their average. Every value is calculated by averaging the sorted sequences' read counts with the next 24 sequences (5-nt, 6-nt, and 7-nt long loops) or 4 sequences (4-nt long loops).

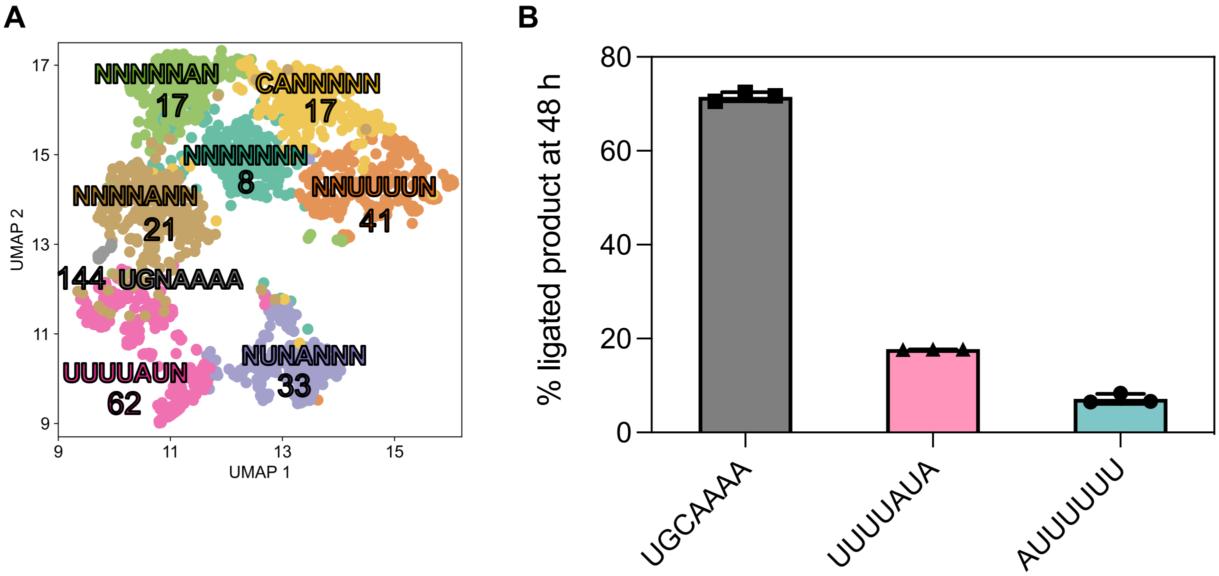


**Fig. S14**. **Cluster analysis of the sequencing data using a K-means algorithm. A:** UMAP projection shows Shapley value space colored according to K-means clusters performed on high-dimensional data. A representative sequence for each cluster is shown here, with an arbitrary threshold of 0.35 for the frequency of a given nucleotide to be considered representative for a given position. Median reads for the cluster are also reported below the representative sequence. **B**: The capture reaction using a representative sequence from each of the top three clusters.

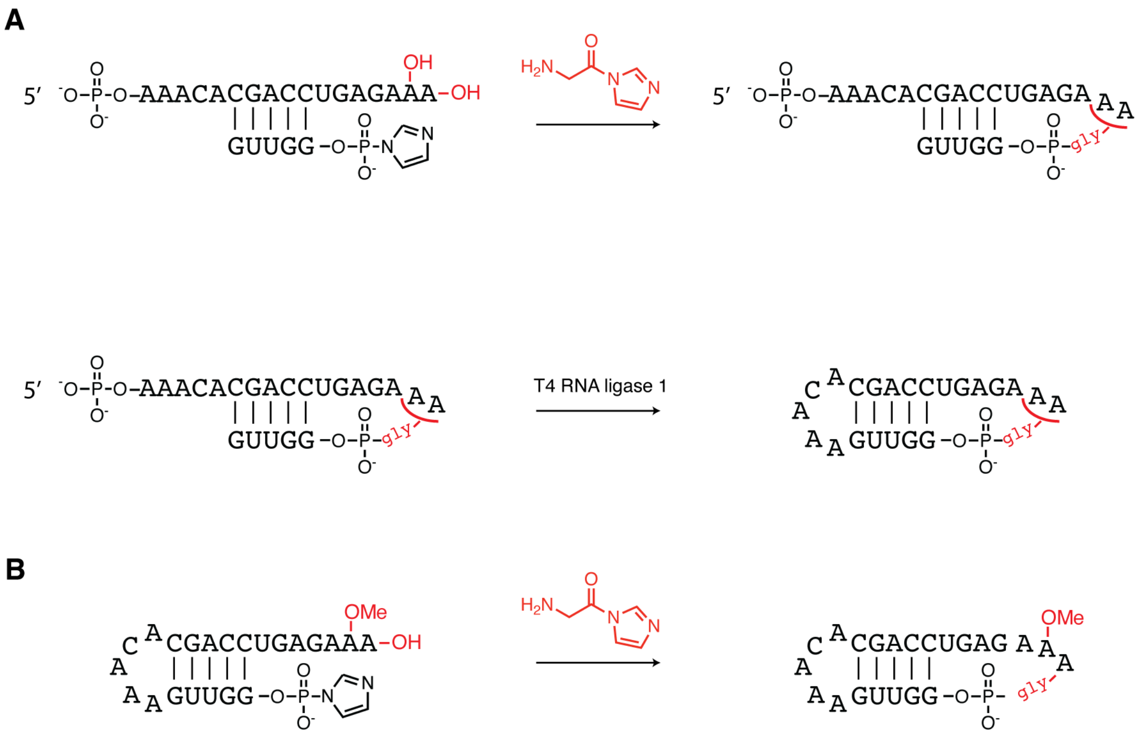


**Fig. S15. Synthesis of the dumbbell shaped RNAs for crystallography. A**: Oligonucleotide with the UGAGAAA-3′ overhang is aminoacylated and ligated to the partially complementary oligonucleotide by incubation with CDI-activated gly. The 5′-AAACA overhang is then closed into a loop with T4 RNA ligase 1. **B**: Oligonucleotide that already has the 5′-AAACA preinstalled during synthesis is closed on the UGAGAmAA-3′ end by incubation with CDI-activated gly.

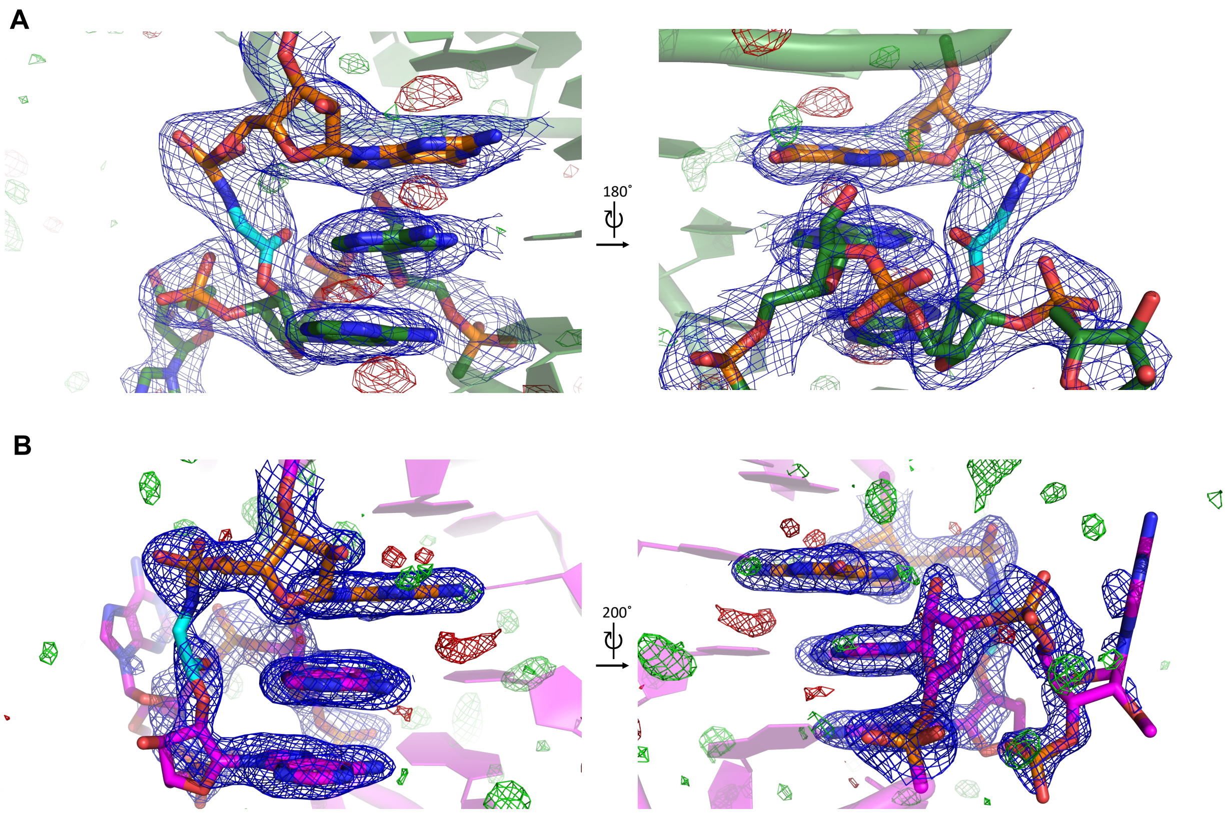


**Fig. S16. Electron density maps corresponding to the glycine linker and residues G1, A20, A21 and A22 in both crystallized constructs**. Blue meshes represent the 2Fo − Fc electron density map at 1.5σ contour level and carve radius 2 Å. Fo − Fc map represented as green mesh at + 3.0 σ and red at − 3.0 σ). **A**: Loop-closed dumbbell RNA bridged by glycine. **B**: Loop-closed A21 2′-OMe dumbbell RNA bridged by glycine.

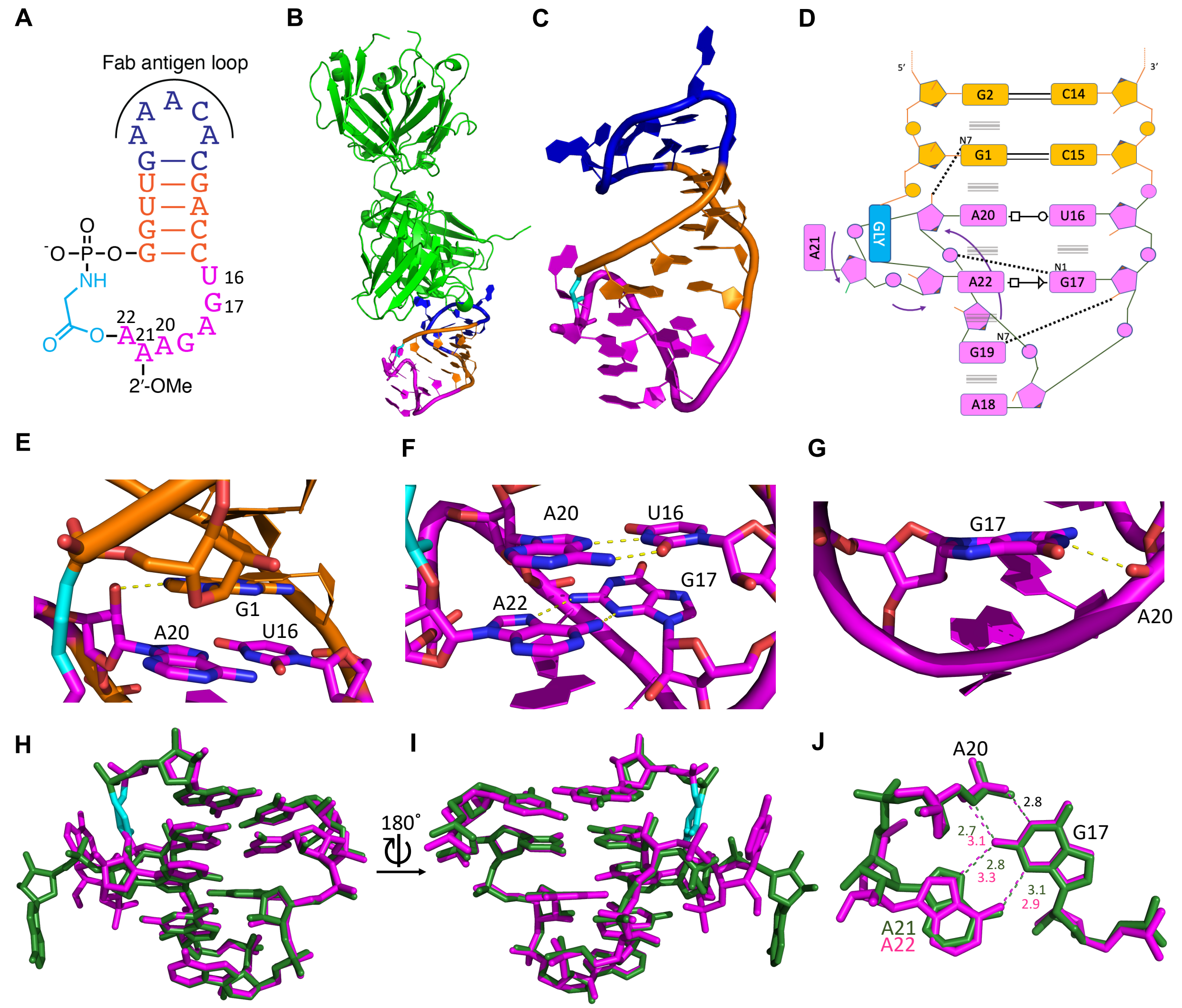


**Fig. S17. Secondary and tertiary structures of the loop-closed A21 2′-OMe dumbbell RNA bridged by glycine determined by crystal structures. A:** Schematic secondary structure of the 2′-OMe modified chimeric dumbbell RNA crystallization construct. **B:** Overall arrangement of loop-closed A21 2′-OMe dumbbell RNA bridged by glycine and Fab BL3-6 in the crystal structure, the closed loop in magenta, glycine linker in cyan. **C:** Tertiary structure of the loop-closed construct. **D:** Secondary structure of the closed loop, interactions are denoted using the Leontis-Westhof nomenclature. **E:** A20:2′OH interacts with G1:N7. **F:** Parallel base pairing interactions in the ligated loop. **G:** G17:N1 interacts with A20:OP2. **H:** Superposition of structures derived from the unmodified dumbbell closed loop (green) and the A21 2′-OMe dumbbell closed loop (magenta), glycine linker in cyan in both structures. **I:** Same as G but rotated 180 in y axis. **J:** Distances in Angstrom and possible interactions (dashed lines) between G17, A20 and intercalating A in the ligated loop crystal structures.

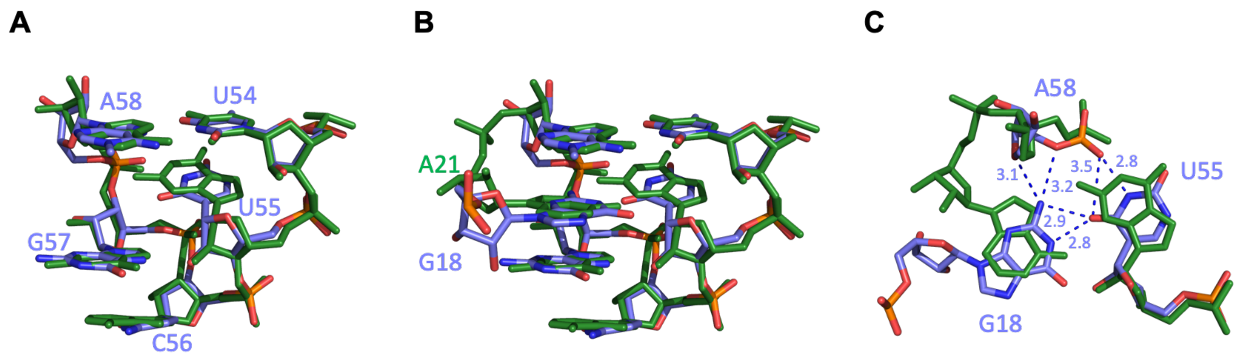


**Fig. S18. The overlay of the tertiary structure of the dumbbell RNA bridged by glycine and the tRNAphe T-loop. A:** Superposition of the closed loop structural motif and the T-loop motif (light blue) from the tRNAphe (PDB 1EHZ) without the intercalating base. **B**: Superposition of the closed loop structural motif and the T-loop motif (light blue) from the tRNAphe (PDB 1EHZ), together with intercalating bases. **C**: Distances and interactions between G18 and U55 and A58 of tRNAphe (PDB 1EHZ) superimposed with the closed glycine-bridged loop (green).

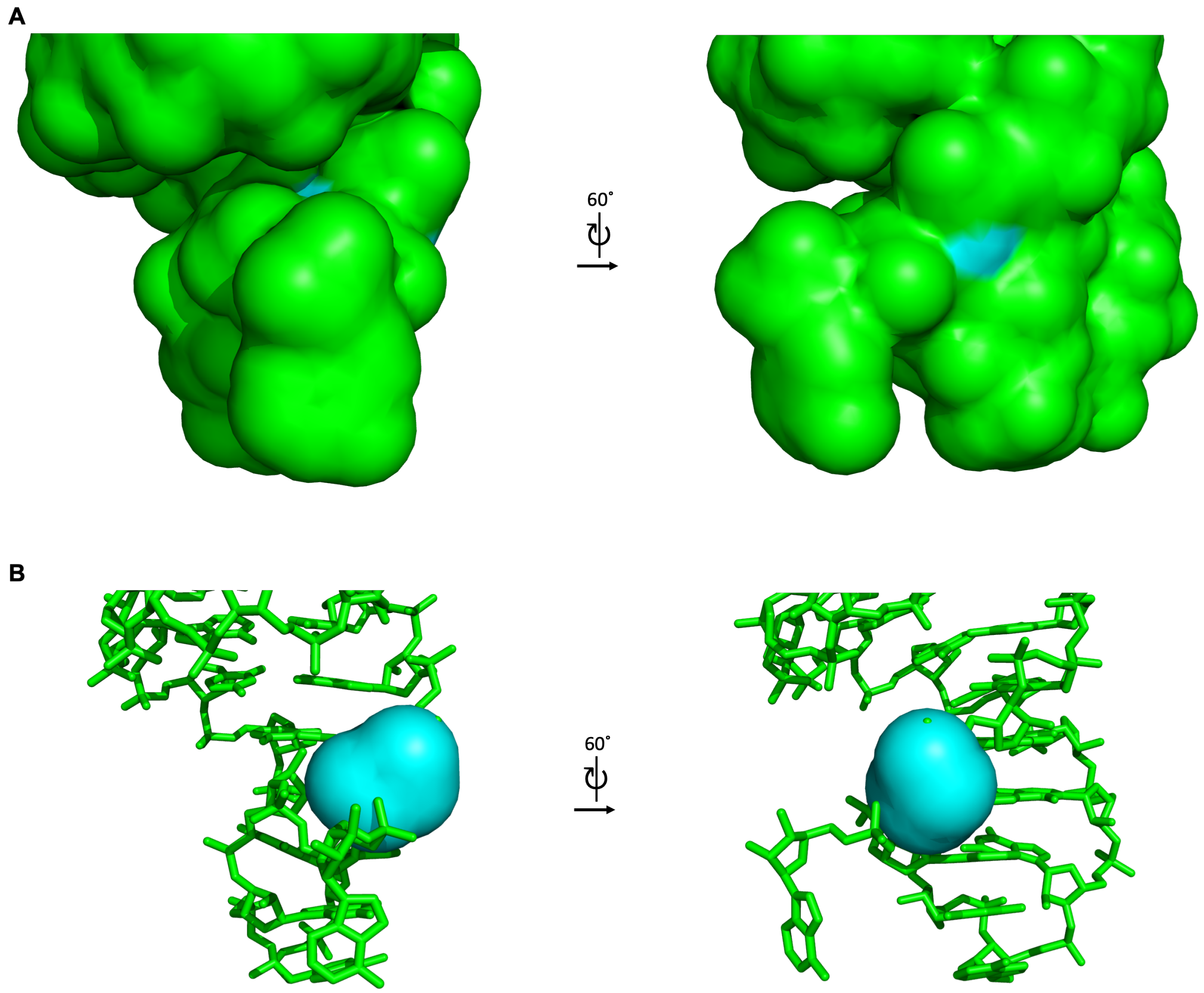


**Fig. S19. Solvent-accessible surface area (SASA) of the glycine-bridged dumbbell RNA.** Figures were generated in PyMOL with the solvent radius probe of 1.4 Å. **A:** SASA presented as solid surface for RNA in green and the glycine bridge in cyan. **B:** SASA presented as solid cyan surface for the glycine bridge only.

**Table S1**.

Stability of the aminoacyl ester linkage in different sequence and structure contexts. Conditions: 200 mM HEPES pH 8, 2.5 mM $MgCl_2$ at 22 °C. [a]value taken from ref. (8) for sequence A4C4 in Table S3. The uncertainty represents the 95 % confidence interval based on three replicates.

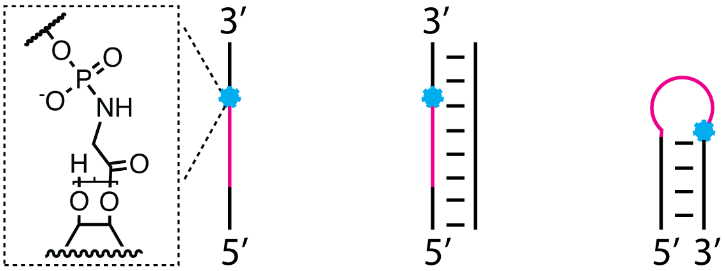


Half life ($h^{-1}$)

| Sequence | ss | ds | loop |
|---|---|---|---|
| AAAA | – | 37 ± 3 | 290 ± 43 |
| UGAGAAA | – | 80 ± 6 | 631 ± 52 |
| GACA[a] | 20 ± 1[a] | 72 ± 2[a] | – |

**Table S2.**
Data collection and refinement statistics. Statistics for the highest-resolution shell are shown in parentheses.

| | **Loop-closed dumbbell RNA bridged by glycine** | **Loop-closed A21 2′-OMe dumbbell RNA bridged by glycine** |
|---|---|---|
| **Wavelength** | 0.979180 | 0.979180 |
| **Resolution range** | 46.07 - 2.07 (2.144 - 2.07) | 41.56 - 1.57 (1.626 - 1.57) |
| **Space group** | P 1 | P 1 21 1 |
| **Unit cell** | 38.514 82.085 92.668 63.953 88.687 88.401 | 39.466 83.126 73.339 90 90.255 90 |
| **Total reflections** | 222414 (22208) | 221740 (20512) |
| **Unique reflections** | 57272 (5776) | 65261 (6414) |
| **Multiplicity** | 3.9 (3.8) | 3.4 (3.2) |
| **Completeness (%)** | 91.14 (91.81) | 96.43 (98.30) |
| **Mean I/sigma(I)** | 6.55 (1.12) | 9.31 (1.48) |
| **Wilson B-factor** | 39.78 | 19.94 |
| **R-merge** | 0.136 (0.9437) | 0.6197 (0.7562) |
| **R-meas** | 0.1576 (1.096) | 0.7276 (0.9121) |
| **R-pim** | 0.07945 (0.5563) | 0.378 (0.5034) |
| **CC1/2** | 0.992 (0.591) | 0.903 (0.488) |
| **CC*** | 0.998 (0.862) | 0.974 (0.81) |
| **Reflections used in refinement** | 56650 (5775) | 63650 (6414) |

| | | |
|---|---|---|
| **Reflections used for R-free** | 2025 (195) | 1993 (199) |
| **R-work** | 0.2211 (0.3020) | 0.1861 (0.2677) |
| **R-free** | 0.2537 (0.3368) | 0.2122 (0.2799) |
| **CC(work)** | 0.953 (0.750) | 0.959 (0.756) |
| **CC(free)** | 0.964 (0.651) | 0.946 (0.602) |
| **RMS(bonds)** | 0.003 | 0.006 |
| **RMS(angles)** | 0.58 | 0.93 |
| **Ramachandran favored (%)** | 97.25 | 97.95 |
| **Ramachandran allowed (%)** | 2.75 | 2.05 |
| **Ramachandran outliers (%)** | 0.00 | 0.00 |
| **Rotamer outliers (%)** | 0.93 | 0.26 |
| **Clashscore** | 3.84 | 3.00 |
| **Average B-factor** | 51.11 | 27.23 |

**Table S3.**

Sequences used in this work. Nucleotides in purple are deoxyribonucleotides, while all others are ribonucleotides. 2-MeImp represents 2-methylimidazole activated phosphate. Imp represents imidazole activated phosphate. Prefix "m" represents 2′-O-methyl nucleotides. "(circ)" indicates the sites of circularization.

| Name | Sequence | Use |
|---|---|---|
| A1 | 5′-fluorescein-AGAGAGCAGACA | Figures 1B, S3 |
| C1 | 5′-Imp-CCCGGCAGCU | Figures 1B, S3 |
| T1 | 5′-AGCUGCCGGGAUGUCUGCUCUCU | Figures 1B, S3 |
| T2 | 5′-AGCUGCCGGGAAUGUCUGCUCUCU | Figure S3 |
| T3 | 5′-AGCUGCCGGGAAAUGUCUGCUCUCU | Figure S3 |
| A2 | 5′-fluorescein-GCAUCCCCA | Figures 1C, S5,6,7,9 |
| C2 | 5′-Imp-AAAGGGUACA | Figures 1C, S5,6,7,9 |
| FX3 | 5′-fluorescein-AGAGAAGCCA | Figure S2A |
| dFX3 | 5′-fluorescein-AGAGAAGCCA | Figure S2B |
| Pc2_m4 | 5′-2-MeImp-CGAAAGUGUACA | Figure S4 |
| Block | 5′-CACTTTCG | Figure S4C |
| A2C2 | 5′-fluorescein-GCAUCCCCA-gly-AAAGGGUACA | Table S1, Figure S8A |
| A2C2duplex | 5′-UGUACCCUUUUGGGGAUGC | Table S1, Figure S8A |
| A2C2revcomp | 5′-GCAUCCCCAAAAGGGUACA | Table S1, Figure S8A |
| A3C3 | 5′-fluorescein-GGACCUGAGAAA-gly-GGUCCCGCAUCCCAGUCUGAGAAA | Table S1, Figure S8B |
| A3C3duplex | 5′-GAGAAUAACGCCAUGUACCAGUCUUUCUCGACUGGGAUGCGGGACCUUUCUCAGGUCC | Table S1, Figure S8B |
| A3C3revcomp | 5′-GGACCUGAGAAAGGUCCCGCAUCCCAGUCUGAGAAAGACUGGUACAUGGCGUUAUUCUC | Table S1, Figure S8B |
| A4C4 | 5′-fluorescein-AGAGAAGAGAGCAGACA-gly-CCCGGCAGCU | Table S1, ref. 8 |
| A5 | 5′-fluorescein-GUUCAGAGUUCUACAGUCCGACGAUCGCGAGGACCNNNN | Figures 2, S10, S11, S13 |

| | | |
|---|---|---|
| A6 | 5′-fluorescein-GUUCAGAGUUCUACAGUCCGACGAUCUUCAGGACCNNNNN | Figures 2, S10, S11, S13 |
| A7 | 5′-fluorescein-GUUCAGAGUUCUACAGUCCGACGAUCCAUCGGACCNNNNNN | Figures 2, S10, S11, S13 |
| A8 | 5′-fluorescein-GUUCAGAGUUCUACAGUCCGACGAUCACAAGGACCNNNNNNN | Figures 2, S10, S11, S13-14 |
| C5 | 5′-Imp-GGUUCAAUAAGAUCGGAAGAGCACACGUCU | Figures 2, S10 |
| RT_3′protector | 5′-AGACGTGTGCTCTTCCGATCT | Figures 2, S10 |
| 5′protector | 5′-GATCGTCGGACTGTAGAACTCTGAAC | Figures 2, S10 |
| 3′-RT_binding | 5′-AppAGATCGGAAGAGCACACGTCT/3ddC | Figures 2, S10 |
| A9 | 5′-fluorescein-GGACCGCGC | Figure 2B |
| A10 | 5′-fluorescein-GGACCGCAA | Figure 2B |
| A11 | 5′-fluorescein-GGACCCGGAU | Figure 2B |
| A12 | 5′-fluorescein-GGACCCGGAA | Figure 2B |
| A13 | 5′-fluorescein-GGACCAUUUUG | Figure 2B |
| A14 | 5′-fluorescein-GGACCAUUUCG | Figure 2B |
| A15 | 5′-fluorescein-GGACCUGUGAAA | Figure 2B |
| A16 | 5′-fluorescein-GGACCUGCGAAA | Figure 2B |
| A17 | 5′-fluorescein-GGACCUGAGAAA | Figures 2, 4, 5, S12 |
| C6 | 5′-Imp-GGUUC | Figures 2B, 4, S12, S14B |
| A56 | 5′-phosphate-AAACACGACCUGAGAAA | Figure S15A |
| C9 | 5′-Imp-GGUUG | Figure S15A |
| chimeric dumbbell RNA | (circ)AAACACGACCUGAGAAA-gly-GGUUG(circ) | Figures 3, S15A, S16A, S17H-J, S18, S19 |
| A57 | 5′-Imp-GGUUGAAACACGACCUGAGAmAA | Figures S15B |
| A21 2′-OMe chimeric dumbbell RNA | (circ)AAACACGACCUGAGAmAA-gly-GGUUG(circ) | Figures S16B, S17 |
| A17– | 5′-fluorescein-GGACCUGAGAA | Figure 4 |
| A18 | 5′-fluorescein-GGACCUGAGAmAA | Figure 4 |
| A19 | 5′-fluorescein-GGACCUIAGAAA | Figure 4 |
| A20 | 5′-fluorescein-GGACCUGAGA2ApA | Figure 4 |
| A21 | 5′-fluorescein-GGACCUUUUAUA | Figure S14B |
| A22 | 5′-fluorescein-GGACCAUUUUUU | Figure S14B |
| Piece 1 (A17) | 5′-fluorescein-GGACCUGAGAAA | Figure 5 |

| | | |
|---|---|---|
| Piece 2 | 5′-Imp-GGUCCCGCAUCCCAUCUGAGAAA | Figure 5 |
| Piece 3 | 5′-Imp-GAUGGUACAUGGCGUUAUUCUC | Figure 5 |
| RNA dFx_S7 | 5′-fluorescein-GGACCUGAGAAAGGUCCCGCAUCCCAUCUGAGAAAGAUGGUACAUGGCGUUAUUCUC | Figure 5C |
| Chimeric dFx_S7 | 5′-fluorescein-GGACCUGAGAAA-gly-GGUCCCGCAUCCCAUCUGAGAAA-gly-GAUGGUACAUGGCGUUAUUCUC | Figure 5C |
| Substrate oligo (A17) | 5′-fluorescein-GGACCUGAGAAA | Figure 5C |
